# An accessible platform to quantify oxygen diffusion in cell-laden hydrogels and its application to alginate-immobilized pancreatic beta cells

**DOI:** 10.1101/2024.09.16.611828

**Authors:** Kurtis S. Champion, Hamid Ebrahimi Orimi, Laurier Gauvin, Jonathan A. Brassard, Berit L. Strand, Richard L. Leask, Corinne A. Hoesli

## Abstract

Hydrogels are commonly used to immobilize mammalian cells, serving various purposes such as providing mechanical cues in three-dimensional cultures and acting as barriers for immunoprotection in transplantation. For instance, islet encapsulation holds promise in delivering insulin-producing cells for diabetes cellular therapy. Cell immobilization, by creating a barrier to bulk fluid motion, leads to diffusion-limited molecular transport and concentration gradients of nutrients such as oxygen being consumed by the immobilized cells. Oxygen mass transport models aid in designing immobilization strategies but rely on input parameters like oxygen diffusivity, often assumed rather than experimentally measured due to limited resources or expertise. We propose an accessible, cost-effective, and easy to operate system to experimentally determine the diffusion coefficient of cell-laden hydrogels, with application tested to alginate-immobilized pancreatic beta cell (MIN6). As compared to water, the oxygen diffusion coefficient was significantly reduced in alginate gels. The oxygen diffusion coefficient was inversely correlated with the dynamic loss modulus for gels with similar chemical composition, and significantly reduced when the alginate concentration was increased from 2% to 5%. The viability of immobilized MIN6 cells was highly dependent both on gel concentration and cell density, as predicted by Thiele modulus and effectiveness factor values calculated from measured oxygen diffusion coefficients. The proposed platform, combining a simple experimental setup and the use of dimensionless numbers, offers a straightforward means to predict maximal diffusion distances in cell immobilization strategies. This platform can be implemented in the rational design of cell encapsulation, immobilized cell culture, and tissue engineering strategies.

## 1. Introduction

Hydrogel-based cell immobilization is used to provide mechanical support or stimuli to cells, to create organized tissue structures, or to create a barrier against harmful external conditions. Hydrogels such as alginate have been used to scale up the culture of mammalian cells [1–3], in 3D bioprinting applications [4–6], as well as to retain or protect grafted cells in transplantation applications [7–11].

One of the earliest applications of alginate imobilization for clinical applications was pancreatic islet encapsulation towards the treatment of type 1 diabetes. By creating a physical barrier between the immobilized insulin-producing cells and components of the immune system such as antibodies and immune cells, certain pathways that lead to graft rejection can be circumvented – a principle termed immunoprotection. With rapid progress towards manufacturing insulin-producing cells from pluripotent stem cells, encapsulation could not only be of value for immunoprotection, but also towards containment and potential retrieval of higher-risk grafts that may contain off-target cells. In all of these applications, the hydrogel can also create a barrier for bulk fluid motion, which can limit molecular transport of key nutrients, growth factors and functional proteins such as insulin, as well as waste products [12–14].

A common challenge in immobilized mammalian cell culture or transplantation is ensuring adequate oxygenation because of its low solubility in water and high rate of consumption, particularly when reaching tissue-like cell densities. Oxygen gradients established by the balance of diffusion through the immobilization material and consumption by cells distributed within can result in cell dysfunction, hypoxia and death – reducing therapeutic potential.

Adequate oxygenation of encapsulated pancreatic islets has been a long-standing engineering challenge [15–18]. In the native pancreas, islets exhibit an intricately structured angioarchitecture, essential for ensuring efficient oxygenation of all cells due to their pronounced aerobic metabolism [18,19]. After isolation from a donor, this native vasculature is disrupted. The cells at the core of avascular islet clusters rely on the external diffusion of oxygen making them much more vulnerable to hypoxia. This issue is exacerbated when encapsulating the cells in a hydrogel due to the addition of another external oxygen diffusion boundary [18]. As compared to microbeads, macroencapsulation - where hundreds of thousands of islets (millions of cells) are delivered in a single device – can exacerbate oxygen gradients due to the local competition for oxygen [16,20,21]. These considerations can be applied not only to islet transplantation devices, but any cell immobilization system where bulk fluid movement is limited.

Several analytical and numerical models have been developed to predict oxygen gradients in islet and other cell encapsulation systems [22–26]. Some of the critical parameters for these types of models include: the cell fraction, the oxygen consumption rate (OCR) of the modelled cell(s), or the oxygen diffusion coefficient of the surrounding medium (ex: cell media, hydrogels, etc.). However, the current limitation is that modelling parameters are most often assumed rather than measured experimentally [27]. Quantifying parameters, such as the oxygen diffusion coefficient is non-trivial but necessary owning to their significant impact on the simulation and variation even within a class of hydrogels.

Existing methods for experimentally measuring the oxygen diffusion coefficient of hydrogels require complex apparatus and specialized equipment. A simple method relies on dissolved oxygen monitoring after previously deoxygenated microbeads in a stirred tank apparatus [28]. This approach offers prompt and dependable evaluations of the oxygen diffusion coefficient, but the absence of cells and gel microstructure observed in beads may not reflect the diffusion coefficient of cell-laden gels of other geometries. More modern methods involve the use of probes like spin probes for electron spin resonance microimaging [29] or optical oxygen dyes and microsensors [30–33]. These methods can provide accurate non-destructive estimates of oxygen diffusion coefficients *in situ* but require specialized equipment which may limit accessibility. Oxygen diffusion coefficients can be estimated from micro-computed tomography (µCT) data [34], which additionally provides information on material structure. This method is versatile but still requires equipment not available in most biology laboratories, and to our knowledge it has not yet been applied to measure oxygen diffusion coefficient through hydrogels such as alginate. Optical oxygen sensors such as microprobes have been broadly applied for point measurements of oxygen concentration values within hydrogels, but the method has not been standardized and approaches vary between studies and encapsulation geometries.

Here, we describe a simple approach to measure the oxygen diffusion coefficient in liquids and hydrogels that allows standardization against measurements acquired for water. The setup can be reproduced at low costs in most wet chemistry and biology laboratories. We demonstrate the application of this system to identify oxygen requirements of alginate-immobilized pancreatic beta cells.

## 2. Materials and Methods

### 2.1. System to measure oxygen diffusion coefficient

The setup to measure oxygen diffusion is shown in **Figure 1A**. A 3D printed cap (see Supplementary, section 1) is attached to a glass Petri dish (316060; Pyrex) and sealed using sealing film (HS234526A, Sigma-Aldrich) to control the oxygen tension at the surface and have negligible oxygen diffusion at the bottom (glass). The positions for the loading port, inlet and outlet hoses were threaded and prepared by installing hose fittings (5117K83, McMaster-Carr), which were sealed with O-rings (5233T368, McMaster-Carr). An oxygen sensing needle probe (OX500PT; Pyroscience) is fixed in place at the center of the system using a universal stand and a positioning mechanism allows for the local oxygen level to be measured various depths (**Figure 1B**). To improve the measurement accuracy, the Petri dish setup is placed in a temperature-controlled environment (**Figure 1A and C**). The gas entering the Petri dish is a mixture of air, N_2_ and CO_2_. CO_2_ was provided through volumetric control of a flowmeter (Cole-Parmer; 03227-04), while larger capacity flowmeters (Cole-Parmer; 03227-12) were used to regulate the amounts of both N_2_ and air. The gases are mixed.

**Figure 1.**
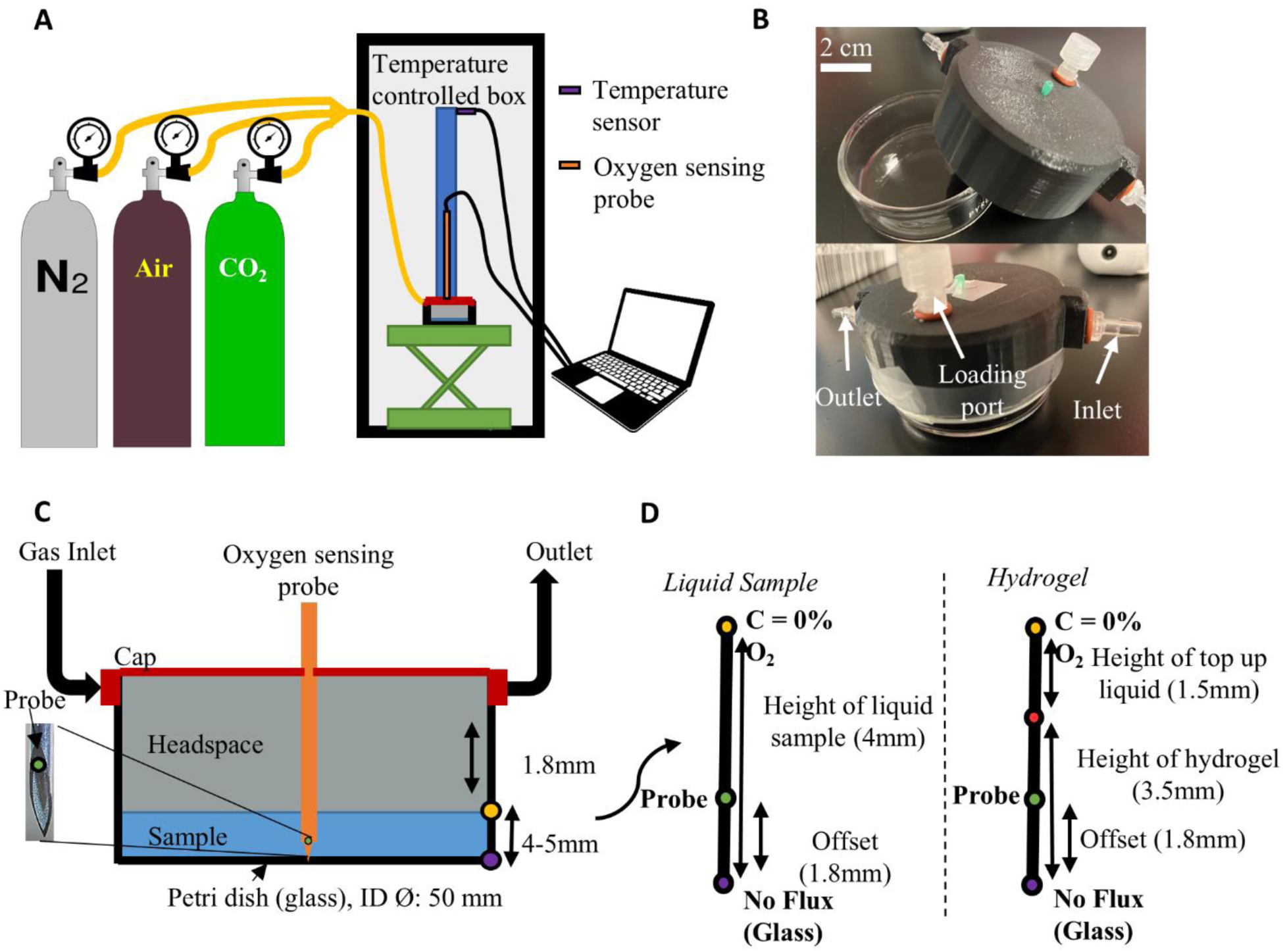
System setup and modelling approach. (A) Exterior system setup. Three blended gases (CO2, N2, Air) are fed into the Petri dish and oxygen is monitored. The setup is fully enclosed in a temperature-controlled environment. (B) Close up images of the Petri dish system when open and sealed closed with sealing film. (C) Technical diagram of cross-sectional view of the Petri dish setup with relevant measurements for modelling. (D) Close up on the sample in the Petri dish. Simplified 1D diffusion model for liquid samples and hydrogels including boundary conditions and modelling relevant measurements. The liquid/hydrogel interface is shown in red.

The oxygen probe is first calibrated using a 2-point calibration in the Pyroscience O2Logger software, the two points are: (i) 0% O_2_ using water with an oxygen scavenger (OXCAL, Pyroscience) and (ii) air-saturated water at a set temperature in the software corresponding to the temperature measured with an alcohol thermometer. The temperature-controlled box was set to 37°C and all liquid or hydrogel samples were equilibrated in a cell culture incubator before testing (37°C; 5% CO_2_). The gas line was purged with 100% nitrogen (N₂) at a flow rate of 300 mL/min for 10 min to eliminate the effects of dead volume in the humidifier and gas lines. Then the flowrate was reduced to 130 mL/min. The OX500PT probe needle was gently placed into the Petri dish sensing port.

A syringe with a 15-gauge blunt-edge needle was used to inject samples into the Petri dish using the loading port (**Figure 1C)**. Initial gas bubbles were removed by flicking the probe. Oxygen measurements were continuously logged in the software with no data smoothing. After waiting 5- 10 min for system stabilization (vibration, temperature, etc.), we attached the N_2_ purged gas line (130 mL/min) to the inlet (**Figure 1C**) and captured the gradual decrease in oxygen over time. An experimental check using water was performed for hydrogel experiments to confirm diffusivity accuracy. **Figure 1D** illustrates the dimensions and positioning of the probe within liquid or hydrogel samples.

### 2.2. Further assessment of oxygen diffusion experimental setup using PDMS-CaO2 slabs

To assess the effectiveness of the oxygen diffusion setup proposed in this study, we quantified the oxygen release rate in slabs composed of polydimethylsiloxane (PDMS) and calcium peroxide (CaO_2_). To prepare the composite, we mixed PDMS base, curing agents (10:1 v/v ratio), and 10% w/v CaO_2_, and then subjected the mixture to vacuum for 20 min to remove air bubbles. The resulting mixture was poured into a glass petri dish and cured in an oven at 50°C for 12 h. Then the slab was removed from the mold and placed at the bottom of the device before placing the cap and sealing.

### 2.3. Computational modelling and governing equations

All modelling was performed in COMSOL Multiphysics 5.6. The modelling was performed in two steps: (1) the boundary condition verification which makes use of the 2D time independent laminar flow module, and (2) the diffusion coefficient numerical model which makes use of the 1D time dependent transport of dilute species module.

### 2.4. Boundary condition verification using computational fluid dynamics

The 2D laminar flow module was used to model the gas flow profile in the headspace between the hydrogel/fluid and the cover of the glass Petri dish. In turn, this model can estimate the water surface velocity and provide an estimate for the Péclet number to determine if the system is diffusion driven. A 2D geometry was used for improved mesh resolution and corresponds to the center plane of the Petri dish where the oxygen probe is located. The simulation assumes incompressible fluid, steady fully developed laminar flow at the Petri dish inlet with the flow rate as set via the gas flow meter, no slip condition at the Petri dish rigid wall, Newtonian fluid, and incompressible fluid (air and liquid region). The boundary layer at the surface of the liquid was taken as an air-liquid interface. The boundary conditions are: (1) normalized laminar flow at the inlet (flowrate per known inlet length scale). (2) zero pressure at outlet (results in gauge pressure profile), and the model is at steady state (time independent). Note that the length scale used to normalize the flowrate was determined by the inner diameter in the inlet tubing, which is 4 mm. An “extremely fine” mesh was used for all calculations (45830 elements; size of 5×10^-4^ to 0.168). The governing equations for solving the velocity and pressure profile were the 2D cartesian time independent Navier-Stokes equation, and the conservation of mass (continuity equation). The shear stress profile was calculated from the product of the shear rate and viscosity (both calculated by COMSOL natively).

The Péclet number (Pe) is the ratio of advective mass transport to diffusive mass transport. It was calculated using the perpendicular surface velocity (u) found from the model at the air-liquid interface boundary layer, the characteristic length (L) which was taken as the height of liquid sample, and the diffusion coefficient (D) (**Equation 1**).

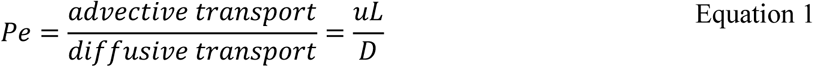

Pe<1 in general suggests that the system is diffusion driven and that recorded data is predominantly a result of diffusion.

### 2.5. 1D time dependent mass transfer model

The 1D transport of dilute species module in COMSOL 5.6 was used to model the oxygen depletion across the center axis of the Petri dish where the oxygen probe is located. The boundary conditions for this line are presented in **Figure 1D**. An extremely fine mesh (45830 elements; size of 5×10^-4^ to 0.168) was used to solve all numerical models, and all reaction terms were ignored since these experiments were acellular. The governing equation for this modelling is Fick’s law (**Equation 2**):

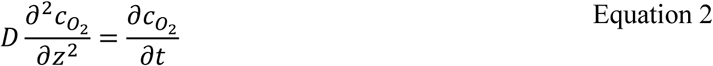

Where z is the length along the center axis where the probe is located.

For a liquid, the top of the line in contact with the gas-controlled boundary condition is assumed to have an oxygen concentration of 0 (C= 0 mol/m^3^) or to follow the transfer function experimentally determined in **Figure 2E** at the appropriate liquid height. The transfer function accounts for the time delay between a step change in oxygen at the inlet and the concentration at the probe location. The opposite end in contact with glass is assumed to have a no flux boundary condition. This assumption is valid since glass has an oxygen diffusion coefficient that is orders of magnitude lower than water and hydrogels. For hydrogel modelling, an additional interface exists between the top up media and hydrogel as shown in **Figure 1D**. Media is assumed to have the same diffusivity as water at 37°C. The probe offset location is determined experimentally and corresponds to where the model is compared to experimental data. The offset distance from the tip of the needle to the beginning of the probe (sheathed within the needle) was measured using a caliper and microscope.

**Figure 2.**
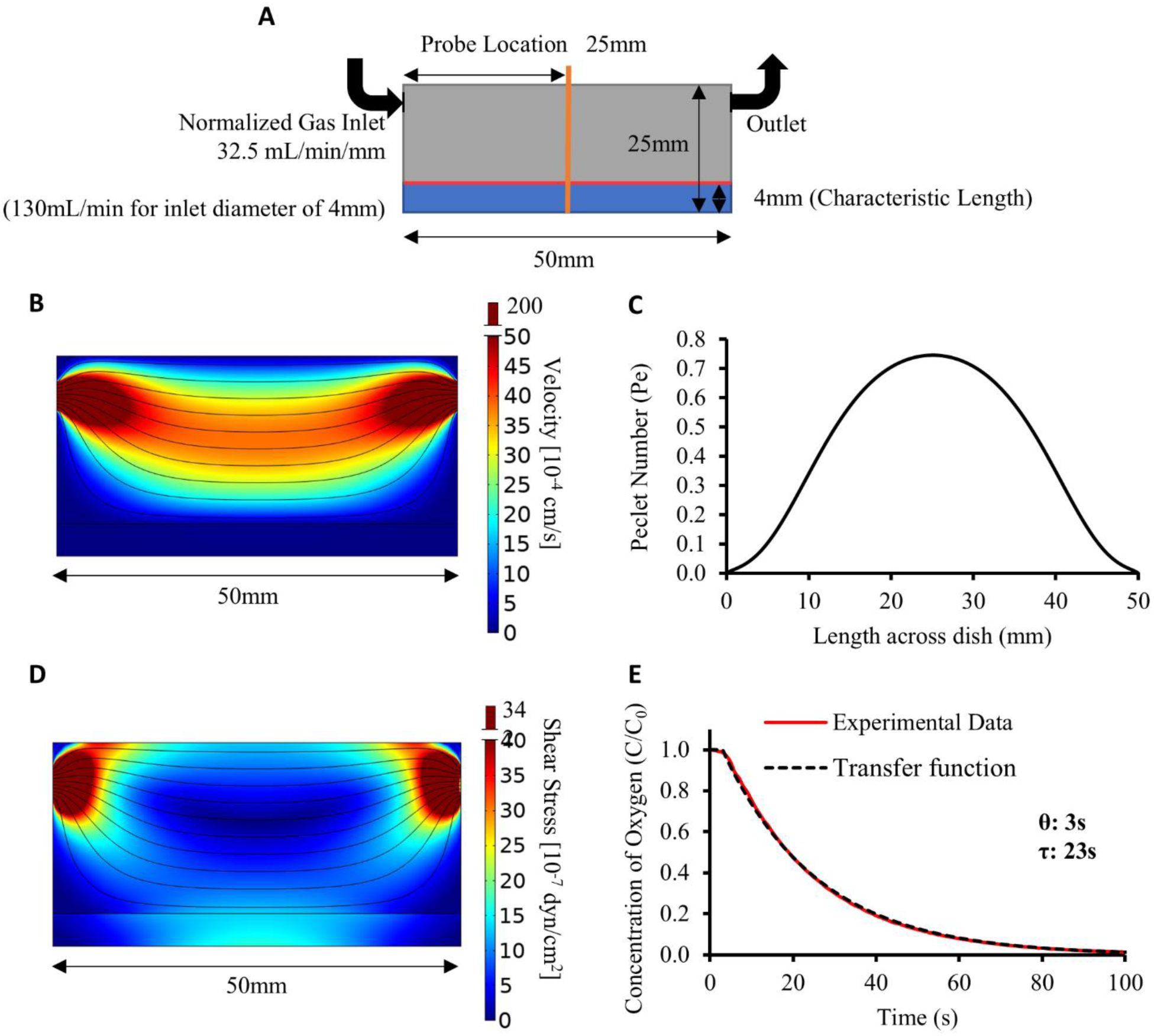
Mass transfer modelling boundary condition justification and assessing advective effects of headspace flow profile. **(A)** 2D geometry simulating the flow profile along the central cross-section of closed Petri dish system with a 50 mm ID Ø, and a gas inlet/outlet port of 4mm ID Ø (D_h_ = 4mm). Includes a sample with a characteristic length of 4mm. The red line illustrates the gas-liquid across the center of the Petri dish. **(B)** The velocity magnitude profile (colour map) and streamlines (black) at the cross-section. **(C)** The Péclet number (Pe) along the gas-liquid interface. Calculated using the perpendicular velocity component (u), the characteristic length (L = 4mm), Pe = uL/D_water_ where D_water_ = ∼3.1×10^-5^ cm^2^/s. Average Pe is 0.48 < 1, max Pe is 0.75< 1. **(D)** The shear stress profile (colour map) and streamlines (black) of the Petri dish cross section. **(E)** Transfer function for the non-dimensional oxygen concentration (C/C_0_) response at the probe location after a step change from 21% O_2_ to 0% O_2_ at the gas inlet. This transfer function is first order with a time delay (θ) of 3 s and a time constant (τ) of 23 s.

### 2.6. Thiele Modulus and Effectiveness Factor

The Thiele modulus (φ) describes the relationship between the reaction rate and the diffusion rate in mass transfer for porous materials like hydrogels. The oxygen consumption in islets was assumed as a 0^th^ order reaction where the cells fixed oxygen consumption rate. 0^th^ order kinetics can be assumed if the ratio between the Michaelis-Menten coefficient (K_m_) and surface concentration (C_s_) approaches 0 (*C*_*s*_ ≫ *K*_*m*_) [35]. This assumption applies to all cells involved in this study, as the oxygen levels never fall below 0.03 mL/m³ (∼25 mmHg), which is higher than the *K*_*m*_of islets (as detailed in the supplementary materials, section 2). Islets have a Michaelis-Menten coefficient of ∼0.44 mmHg for oxygen consumption [22,36] which is much less than the surface oxygen tension of at least 40 mmHg [37] observed for immobilized islet systems when transplanted into an environment where the device surface is exposed to oxygen tensions at the lower end of venous values.

The Thiele modulus for a simple 0^th^ order system was derived from [38]. It was calculated with a constant reaction rate (k_v_) equal to the cellular oxygen consumption rate (OCR) normalized by cell fraction (X), and surface oxygen concentration (C_s_) and is presented as the following (**Equation 3**):

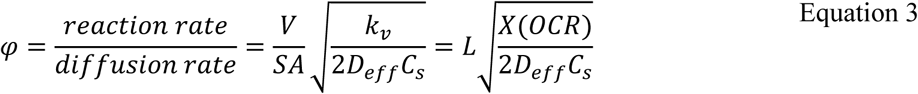

Where V is the volume, SA the surface area and L is the thickness.

The effectiveness factor was calculated from literature for a slab geometry with a 0^th^ order reaction [39]. Note that for this derivation, φ_crit_ is taken as √2 [39] (**Equation 4**).

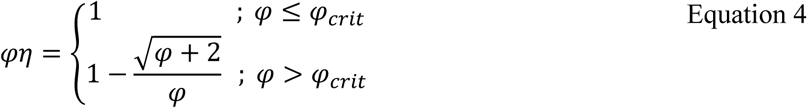

### 2.7. Adherent cell culture

Mouse insulinoma 6 (MIN6) cells (generously gifted from Dr. Jun-ichi Miyazaki- Osaka University [40]) in complete medium were seeded in tissue culture treated polystyrene flasks until 80-90% confluency. The complete medium consisted of Dulbecco’s Modified Eagle medium (DMEM; Gibco #10313-021, Thermo Fisher Scientific) supplemented with 10% fetal bovine serum (FBS; HyClone #SH3039602, Fisher Scientific), 1% L-glutamine (Gibco #25030-081, Thermo Fisher Scientific), 1% penicillin/streptomycin (Gibco #15140-122, Thermo Fisher Scientific), and 0.1% 2-mercaptoethanol (Fisher Chemical #O3446I, Fisher Scientific). Changes were performed every 48-72 h. An incubator at 37°C and 5% CO_2_ was used to culture the cells. Passaging was performed using 1X TrypLE Express Enzyme (Gibco #12605-028, Thermo Fisher Scientific). Experiments were conducted at passage number between 28 and 40.

### 2.8. Production of alginate slabs

Alginate slabs were produced using internal gelation. The two types of alginate used in this work were Manugel GHB (IFF Nutrition & Biosciences (formerly FMC Biopolymer) through DuPont) and Protanal 10/60 LF (IFF Nutrition & Biosciences (formerly FMC Biopolymer) through DuPont). A stock solution of either 2.5% w/v Manugel GHB, 2.5% w/v Protanal, or 6.25% w/v Manugel GHB was prepared in pH 7.4 HEPES-buffered saline (170 mM NaCl; 10mM HEPES). The alginate stock solution was autoclaved at 121°C for 30 min; 100 mL of the stock solution was autoclaved each time. A second stock solution of 0.5 M calcium carbonate (CaCO_3_; Avantor #1301-01, VWR) was also prepared in pH 7.4 HEPES-buffered saline and sonicated for 15 min to disperse calcium carbonate particles. Thereafter, this solution was autoclaved at 121°C for 30 min. Glucono-δ-lactone (GDL; Sigma-Aldrich #G4750, MilliporeSigma) was dissolved in HEPES-buffered saline and sterilized with a 0.2-µm nylon syringe filter (Fisherbrand™ #09-719C, Fisher Scientific) immediately before use.

To prepare a slab with a height of at least 4 mm with a 50 mm diameter, at least 5.6 mL of alginate solution is necessary. For all experiments, 10 mL of alginate hydrogel mixture was made. To do so, 0.6 mL of the 0.5 M CaCO_3_ solution was added into 8 mL of 2.5 % alginate or 6.25 % alginate and mixed using a sterile spatula in a 10 mL syringe with the plunger removed and the bottom capped. A 1 mL solution of cell culture media containing trypsinised MIN6 cells that will result in the desired final cell concentration was then added to the syringe and mixed using the spatula.

Finally, 0.6 mL of the GDL solution was added to the syringe. The resulting solution is ∼2% or ∼5 % w/v alginate stock, 30 mM CaCO_3_, and 60 mM GDL. The syringe plunger was then reintroduced, while preventing any alginate solution from leaking out. The solution, 7.9 mL, was injected and evenly spread into glass Petri dish (Ø 50 mm). Note that 7.9 mL corresponds to a 50 mm cylinder with a 4 mm height (characteristic length used for Pe and modelling). Hydrogel samples include a small amount of top up media to prevent the gel from drying out.

The slabs were left to gel for between 30-40 min (until visually solid) and topped up with 4 mL of cell culture media to prevent the slab from drying out. The Petri dish was then sealed with a piece of parafilm that was previously soaked in ethanol. The slabs were placed in an incubator at 37°C and 5% CO_2_ for 16-24h.

### 2.9. Live/dead staining & image acquisition

After the slabs underwent 16-24h of incubation, a very thin vertical slice of the center of the slab was cut using a razor blade and stained using a live/dead solution with final concentrations consisting of 18.9 µg/mL propidium iodide (Fisher Scientific; P1304MP) and 1.1 µg/mL Calcein AM (Fisher Scientific; C1430) both in pH 7.4 HEPES-buffered saline at room temperature. Images were acquired using an IX81 Olympus Microscope at 10X magnification on the full chip setting using a FITC filter cube (Ex: 482/35 | Em: 536/40) and a Texas Red filter cube (Ex: 525/40 | Em: 585/40). The exposure time used for all imaging was 100ms.

### 2.10. Image analysis for live/dead staining

Images were stitched in a grid-wise format using a stage file generated in MATLAB [41], a step size of 850 μm in the X and Y direction. Live/dead fluorescence analysis and live MIN6 cells depth analysis was performed in FIJI (ImageJ). Acquired images were cropped from 2048×2048 pixels to 1550×1550 pixels and stitched column-by-column, up-and-right, with an overlap of 18% using the built-in stitching function [42]. FITC and Texas Red images were merged where FITC was chosen to be green and Texas Red which was red. Live MIN6 cells depth analysis was performed by setting a calibrated scale of 1300 pixels:850 μm and by visually measuring the live height (the green portion) and total slab height at the top, middle, and bottom of the slab using the line tool in ImageJ.

### 2.11. Alginate characterization

Alginate composition was characterized by ^1^H-NMR as described previously [43–45]. Briefly, the alginate was degraded by acid hydrolysis and lyophilized. 5-10 mg dry sample was dissolved in 600 μl D2O (99.9%) and added 5 μl 3-(Trimethylsilyl) propionic 2,2,3,3-d4 acid (TSP, Sigma Aldrich) as an internal standard. 1D spectra was recorded at 90°C on a 600 MHz spectrometer (Bruker BioSpin AG, Fällanden, Switzerland). The spectra were recorded using TopSpin 1.3 or 2.1 software and processed and analyzed with TopSpin 3.0 software (Bruker BioSpin). The anomeric protons in alginates have chemical shifts in the region 4.3–5.3 ppm, and fractions of monomers (F_G_, F_M_), dimers (F_GG_, F_MM_, F_GM,MG_), trimers (F_GGG_, F_MGM_, F_GGM,MGG_) and minimum G-block length (N**G>1)** was calculated from the intensity profiles in this region as described previously [43–45].

The weight average molecular weight (M_W_), number average molecular weight (M_n_) and polydispersity index (M_w_/M_n_) of the alginate samples was analyzed by Size Exclusion Chromatography with Multi Angle Light Scattering (SEC-MALS) [46]. Samples and standards were dissolved in the mobile phase (0.05M Na_2_SO_4_ with 0.01M EDTA, pH 6.0) and filtered (0.8 μm). Astra software v. 6.1 (Wyatt, USA) was used to collect and process the data obtained from the light scattering and the differential refractometer, using a refractive index increment (at constant chemical potential), (dn/dc)_μ_ of 0.150 ml/g [46].

### 2.12. Rheology

A modular compact rheometer (MCR 302, Anton Paar) was used to measure the storage modulus (G*’*) and loss modulus (G*"*) of hydrogels composed of 5% (w/v) Manugel, 2% (w/v) Manugel, and 2% (w/v) Protanal. The hydrogels were fabricated into slabs approximately 25 mm in diameter and 2 mm in thickness using internal gelation approach. Hydrogel slabs were incubated in Dulbecco’s Modified Eagle Medium (DMEM) at 37°C with 5% CO_2_ and humidity for 24 h.

To study the shear stress experienced by cells, the hydrogel slabs were compressed with a constant axial force (0.2N) to prevent slipping and subjected to an oscillation strain of 0.5%, while varying the applied angular frequency between 0.1 and 100 rad/s. Furthermore, the hydrogel’s elasticity was investigated under an amplitude sweep by keeping the strain rate constant at 10 s^-1^ and varying the oscillation amplitude from 1% to 100%.

### 2.13. Statistical analyses

All averages and standard deviations were computed using Microsoft Excel. The data was plotted in either Excel or in GraphPad Prism (9.1.2). All data underwent normality checks in GraphPad utilizing the Shapiro-Wilk test. To evaluate differences between experimental groups, a one-way ANOVA was performed, followed by the Tukey test to compare groups. Oxygen diffusion coefficients were determined by minimizing the mean absolute percent error (MAPE) of measured oxygen concentration profiles as a function of time to numerical predictions as determined by the 1D time dependent model.

## 3. Results

A straightforward way to measure the oxygen diffusion was developed in this work. Using an oxygen probe and conventional laboratory equipment, the oxygen diffusivity of various hydrogel formulations was measured. One of the main assumptions in this measurement method is that the probe is measuring a system that is diffusion dominant with negligible advective effects. To verify negligible advection, a computational fluid dynamics (CFD) model of the headspace was developed to determine the Péclet number (ratio of advection to diffusion).

### 3.1. Numerical and experimental validation of the platform

To assess whether the geometry and gas flow rates selected would lead mainly to diffusion-driven transport, the gas velocity and shear stress profiles were modeled numerically (**Figure 2**). The flow profile experienced at the sample interface where the probe is inserted was assumed to be the same as the flow profile across the center plane of the Petri dish. While this assumption neglects edge and curvature effects, this will over-estimate advection effects since the velocity at the center of the 2D plane would be higher than in a 3D Petri dish where the cross-sectional area increases with distance from the gas inlet.

With the computed model, the perpendicular velocity at the air-liquid interface (**Figure 2B**) can be used to calculate the Pe number across the Petri dish (shown in **Figure 2C**). The average Pe was 0.48 and the maximum was 0.75, both of which are less than 1 indicating the system is diffusion dominated. To verify the assumption in the mass transfer model that the surface of the hydrogel or water at 0% O_2_, the boundary should decrease rapidly to 0% upon a step change. In **Figure 2E**, the experimental transfer function for the system is presented demonstrating that the boundary goes from 21% to 0% in roughly 100s. Since experiments occur on the timescale of 1- 10h, this would suggest that the 100s transitory period from 21% to 0% is negligible.

To validate the system experimentally, the diffusion coefficient of water at 37 °C was estimated using the setup was compared to literature values which range from 2.9×10^-5^ -3.3×10^-5^ cm^2^s^-1^ [27,47–50]. To estimate the diffusion coefficient from experimental measurements, the oxygen concentration response measured by the probe (center, bottom of the Petri dish) was compared to the unidimensional time-dependent mass transport model (**Figure 3**) to extract the diffusion coefficient providing the best fit. The response observed experimentally showed faster initial response which may reflect some advection effects, but most of the deviations remained within experimental confidence intervals. The value obtained was 3.1×10^-5^ cm^2^s^-1^, which is within the reported range. Water diffusion coefficient values were acquired before each alginate measurement, and results were reproducible.

**Figure 3.**
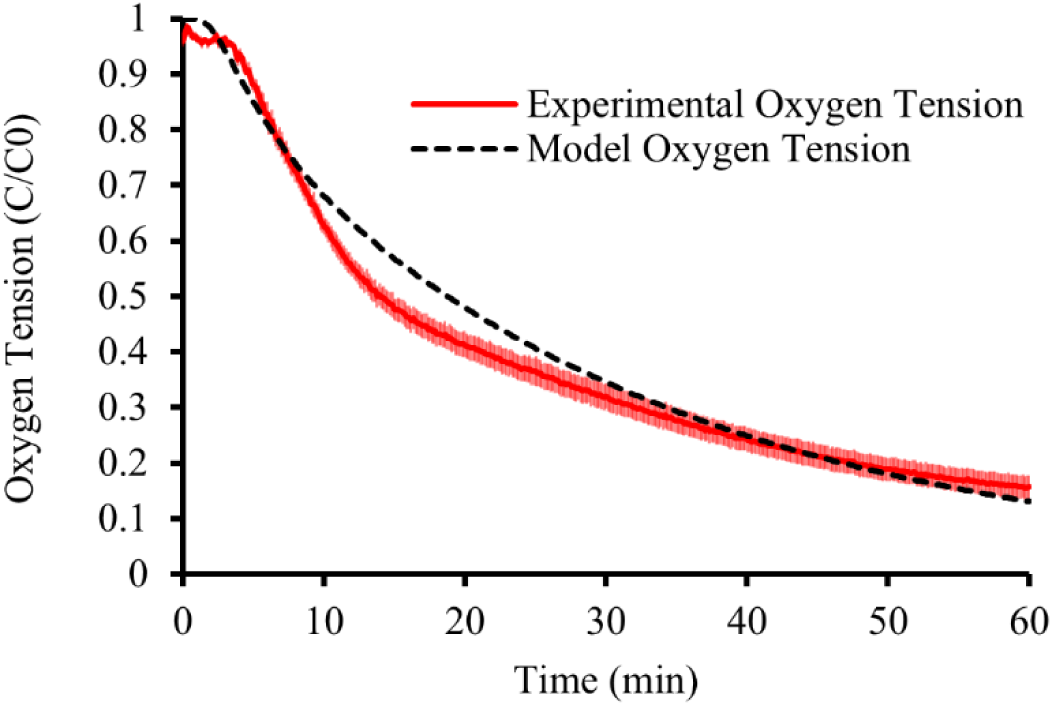
System validation using water at 37°C, Experimental non-dimensional concentration data (red) of oxygen depletion in air-saturated water (n=7). Numerical model with diffusion coefficient of water at 37°C (D∼= ∼3.1×10^-5^ cm^2^s^-1^) (confidence interval represents the standard error of the mean, N=7)

To assess the experimental system’s capacity, a combination of PDMS and CaO_2_ was utilized as an oxygen producer. This method enabled the quantification of the amount of oxygen generated during oxygen purging in the setup. The oxygen release profile of the CaO2/PDMS beads is presented in the supplementary materials, alongside the oxygen release profile of the PDMS beads (Section 3).

### 3.2. Alginate oxygen diffusion coefficient measurements

The system was then used to measure the experimental oxygen diffusion coefficients of different formulations of alginate. As for water, the fitted and experimental oxygen concentration response after a step decrease in oxygen at the system inlet followed similar but not identical trends. Experimental responses were initially delayed as compared to model fit, particularly for the higher-concentration alginate condition. This is characteristic of systems with Pe numbers above 1 [51,52], and could be due to radial diffusion effects neglected by the model that would be more prominent in gels with lower diffusion rates. The gel with the slowest response was the 5% Manugel formulation, followed by the 2% Manugel and the 2% Protanal gel, as also reflected by the fitted diffusion coefficient values (**Figure 4** and **Figure 5**). There was no significant difference between water and 2% Protanal. Although the diffusion coefficient differences between 5% and 2% Manugel did not reach statistical significance, the faster trend observed in the raw data may indicate an effect of the polymer concentration – as would be expected for more tortuous diffusion paths created by polymer chains [53,54]. However, this effect was not as important as the difference between polymer sources, highlighting the limitation of using literature values when estimating alginate diffusion coefficients.

**Figure 4.**
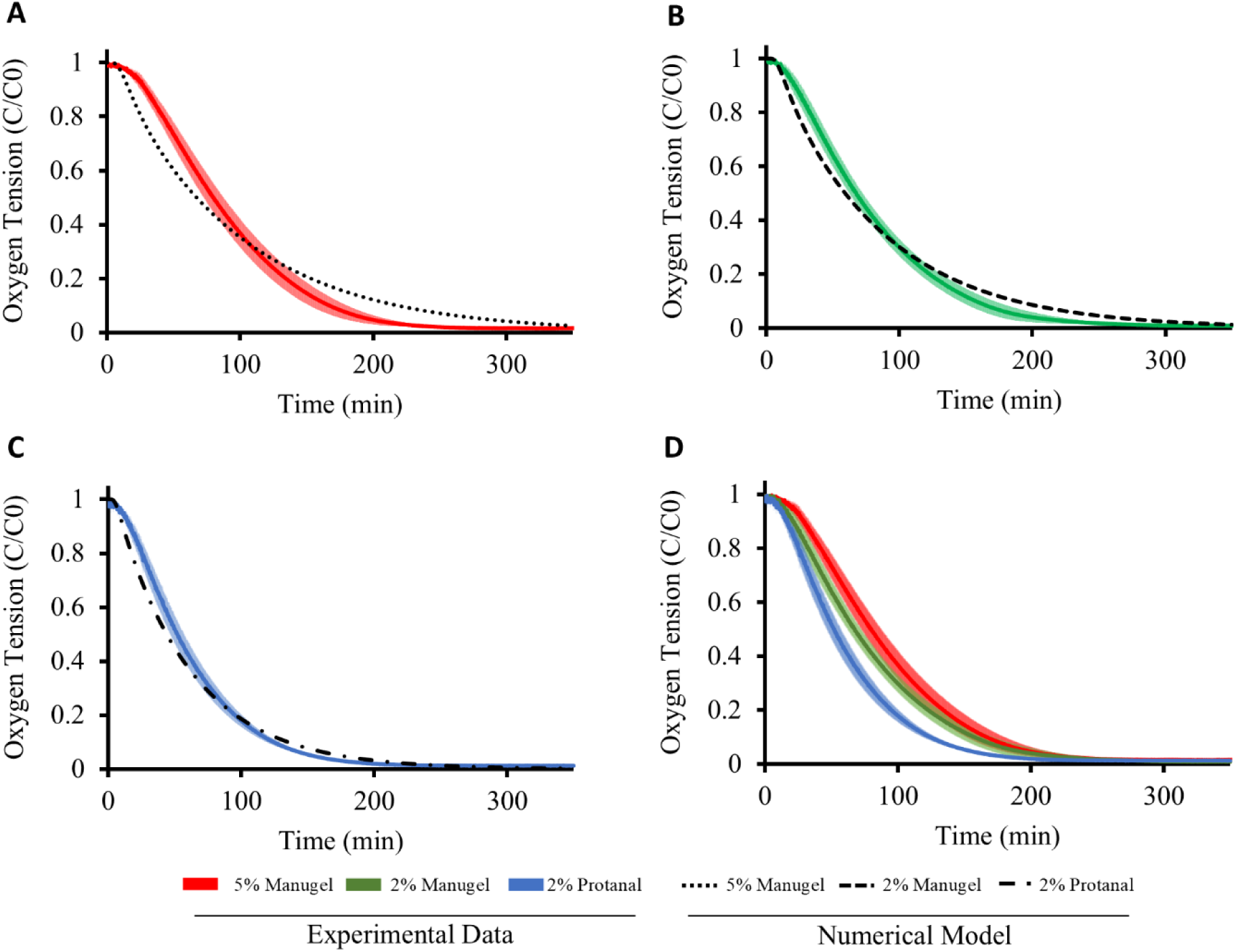
Diffusion coefficient of various alginate compositions at 37°C. (A) Experimental non-dimensional concentration data (red) of oxygen depletion in 5% w/v Manugel alginate in HEPES buffer (n=3) with numerical model corresponding to D= 1.1×10-5 cm2s-1. (B) Experimental non-dimensional concentration data (green) of oxygen depletion in 2% w/v Manugel alginate in HEPES buffer (n=3) with numerical model corresponding to D= 1.5×10-5 cm2s-1. (C) Experimental non-dimensional concentration data (blue) of oxygen depletion in 2% w/v Protanal alginate in HEPES buffer (n=3) with numerical model corresponding to D= 2.7×10-5 cm2s-1. (D) Overlay of experimental data of all alginate formulations tested. (confidence interval represents the standard error of the mean, N=3 to 4).

**Figure 5.**
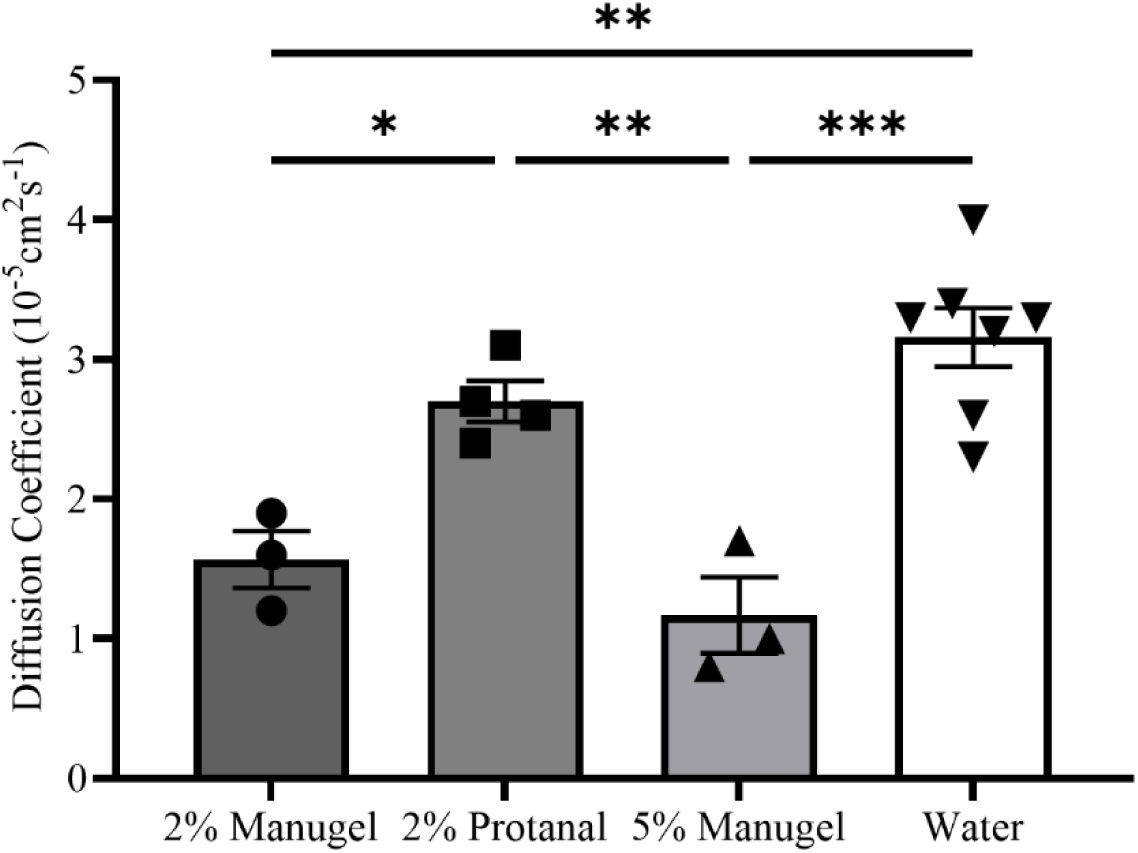
Comparative analysis of diffusion coefficient of various alginate compositions at 37°C, Diffusion coefficients were obtained by minimizing the mean average percent error between experimental data and the unidimensional time-dependent mass transport model. Experimental replicates: n=3 for 2% Manugel and 5% Manugel, n=4 for 2% Protanal, and n=7 for water. Significance was assigned as followed: * p≤0.05, ** p≤0.01, *** p≤0.001. (error bar represents the standard error of the mean, N=3 to 4)

To determine whether the observed differences in oxygen diffusion coefficient between Manugel and Protanal may be due to differences in guluronic acid (G) content, higher molecular weight, or higher number of G-blocks, we assessed the chemical composition of the gels. Surprisingly, both gel types showed very similar composition, indicating that more fine differences – such as regional differences within the chains – could be at play (**Table 1**).

**Table 1.** Molecular Weight, M/G Ratio, and Chain Analysis of Alginate Variants Manugel GHB and Protanal 10/60 LF, in Powder and Autoclaved Forms.

| Alginate | Molecular Weight |  |  | M/G Ratio | Mannuronic Acid (M) / Guluronic Acid (G) Chain Analysis |  |  |  |  |  |  |  |  |
| --- | --- | --- | --- | --- | --- | --- | --- | --- | --- | --- | --- | --- | --- |
|  | M<sub>w</sub> (kDa) | M<sub>n</sub> (kDa) | M<sub>w</sub>/M<sub>n</sub> |  | F<sub>G</sub> | F<sub>M</sub> | F<sub>GG</sub> | F<sub>GM</sub>,<br>M<sub>G</sub> | F<sub>MM</sub> | F<sub>GGM</sub>,<br>M<sub>GG</sub> | F<sub>MGM</sub> | F<sub>GGG</sub> | N<sub>G>1</sub> |
| Manugel GHB (powder) | 176 | 73 | 2.4 | 0.47 | 0.68 | 0.32 | 0.57 | 0.11 | 0.21 | 0.041 | 0.071 | 0.53 | 15 |
| Autoclaved Manugel GHB (2% w/v) | 90 | 43 | 2.1 |  |  |  |  |  |  |  |  |  |  |
| Protanal 10/60 LF (powder) | 143 | 64 | 2.2 | 0.49 | 0.67 | 0.33 | 0.55 | 0.11 | 0.22 | 0.039 | 0.078 | 0.52 | 15 |
| Autoclaved Protanal 10/60 LF (2% w/v) | 82 | 41 | 2.0 |  |  |  |  |  |  |  |  |  |  |

Qualitatively, we had observed that the Protanal gels, at the same concentration, appeared softer than the Manugel gels. We therefore hypothesized that rheological features may correlate with the oxygen diffusion coefficient values. We found that the loss modulus was lowest for 2% Protanal, followed by 2% Manugel and 5% Manugel (**Figure 6**). This trend was consistent with the oxygen diffusion rates, with lower loss moduli correlating with higher oxygen diffusion rates (and thereby estimated oxygen diffusion coefficients). The loss modulus indicates the viscosity of the hydrogel, with higher values indicating greater viscosity. Furthermore, viscosity shows a direct proportionality to the diffusion rate. This explains the correlation observed between the loss modulus and diffusion rate in this study. Slight difference was found between the storage modulus values of the three hydrogels. To assess the significant differences in elasticity among the hydrogels that explain their varying softness, a rheological test was conducted using varying oscillation amplitudes. This test revealed differences in the amount of stored energy (detailed in the supplementary materials, section 4).

**Figure 6.**
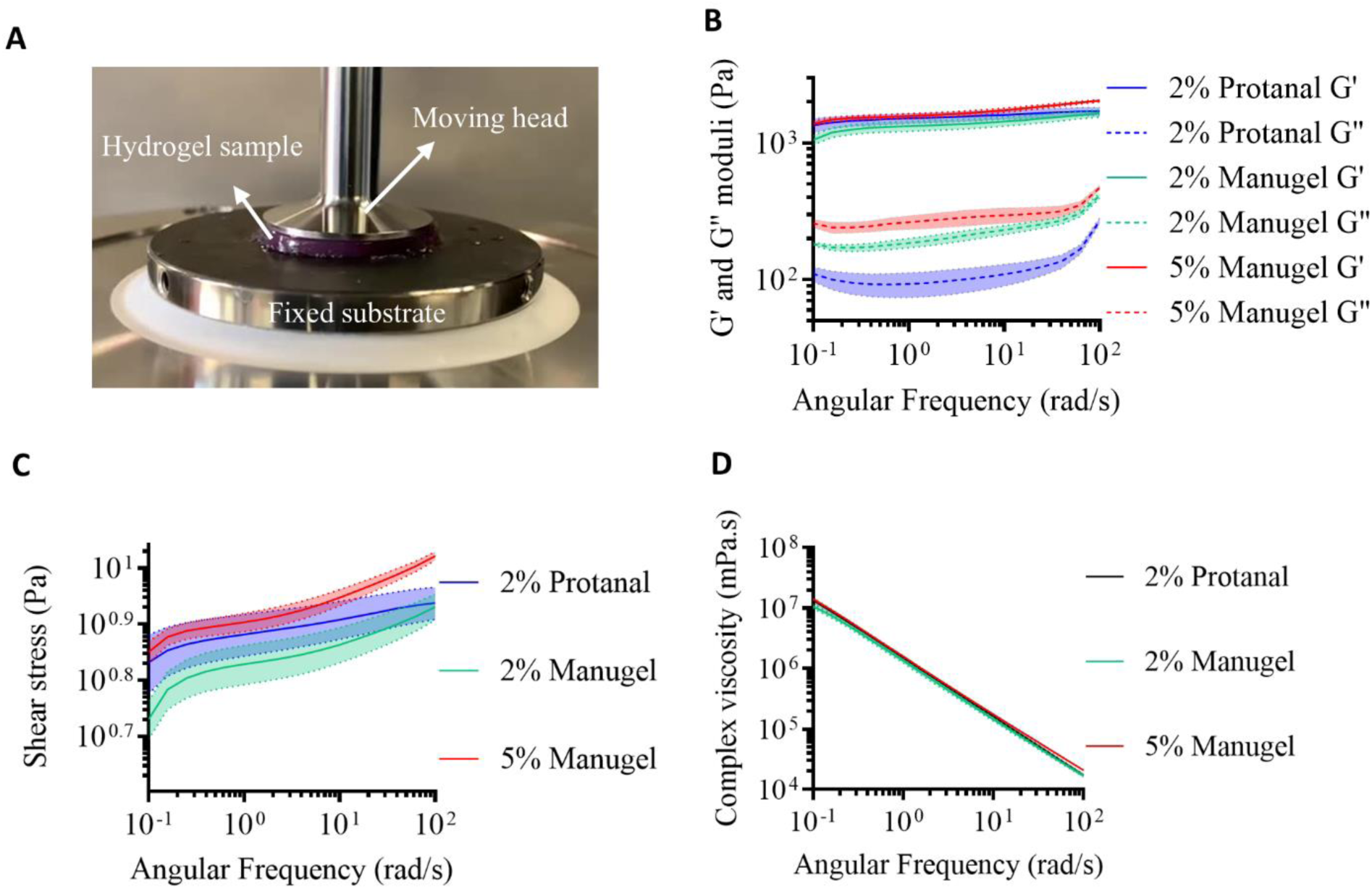
Frequency sweep experiment in rheology of hydrogels at room temperature. **(A)** The hydrogel sample placed under the rheology machine. **(B)** Experimental storage and loss modulus versus oscillation frequency (0.1 to 100 Hz). **(C)** Experimental shear stress versus oscillation frequency (0.1 to 100 Hz). **(D)** Experimental complex viscosity versus oscillation frequency (0.1 to 100 Hz) for 5% w/v Manugel alginate (red zone), 2% w/v Manugel alginate (green zone) and 2% w/v Protanal alginate (blue zone). (confidence interval represents the standard error of the mean, N=3)

### 3.3. 24h MIN6 cell viability and theoretical cell fractions based on Thiele modulus

Since the experiment was done in a simple 1D slab geometry, the effective length (L) is the height of the gel (3.5mm as per **Figure 1D**). The diffusion (D_eff_) was taken as a weighted average using the cell fraction between the diffusion coefficient of tissue (1.24×10^-5^ cm^2^ s^-1^ [23,49]) and the diffusion coefficient of the material as per **Figure 5**. The OCR used for MIN6 cells is 0.129 mol·m^-3^ s^-1^ [16] and 0.034 mol·m^-3^s^-1^ [16,22,23,25] for human islets, note that the MIN6 cells used are from the same cell stock used in [16]. The oxygen surface tension will not always be exactly 18.6% O_2_ since there is still a layer of top up media approximately 1.5 mm high that oxygen must diffuse through before reaching the top of the slab (as per **Figure 1D**). To determine the surface oxygen levels of the slab, a separate numerical model (as detailed in the supplementary materials) was made to develop a relationship between the surface oxygen tension (C_s_) and the cell fraction (X) for a given OCR (MIN6 cells or islets) and alginate diffusion coefficient. Finally, a theoretical cell concentration (X) was iterated to yield an effectiveness factor of 0.9 and 1, this gives us a rough approximation of the maximum cellular density for a given geometry and encapsulation material (**Table 2**).

**Table 2.** Theoretical maximum cell fraction of MIN6 cells and islets to achieve an effectiveness of 0.9 (η = 0.9) and 1 (η = 1)

| Hydrogel | Maximum MIN6 cell fraction<br>(million cells/mL) |  | Maximum islet cell fraction<br>(million cells/mL) |  |
| --- | --- | --- | --- | --- |
|  | For effective factor of 0.9 | For effective factor of 1 | For effective factor of 0.9 | For effective factor of 1 |
| 2% Manugel | 1.10 | 0.92 | 3.5 | 3.05 |
| 2% Protanal | 1.60 | 1.35 | 5.3 | 4.65 |
| 5% Manugel | 8.50 | 7.1 | 2.8 | 2.40 |

To determine the relationship between diffusion and cell behaviour, the viability of MIN6 cells in a 1D diffusion slab geometry was evaluated. The experimental viability of MIN6 cells at two different cell fractions is presented in **Figure 7**.

**Figure 7.**
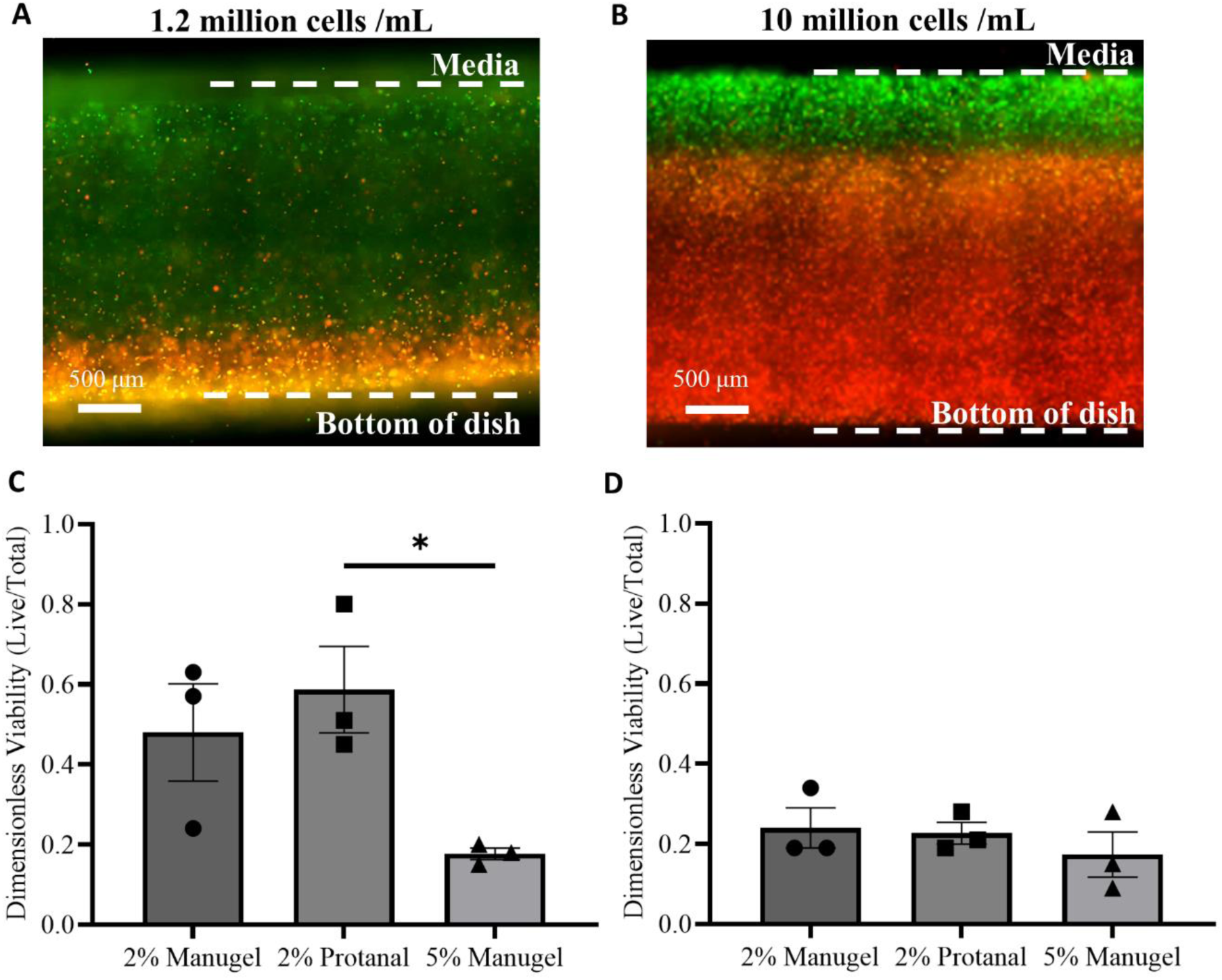
Live/dead analysis of MIN6 cells at 1.2 million cells per mL and 10 million cells/mL, Example image of 2% protanal live/dead cross-section at 1.2 million MIN6 cells/mL. Green indicates live cells (FITC; Ex: 482/35 | Em: 536/40), red indicates dead cells (Texas Red; Ex: 525/40 | Em: 585/40) after 16-24h of incubation (B) Example image of 2% protanal live/dead cross-section at 10 million MIN6 cells/mL. (C) Dimensionless viability of MIN6 cells at a concentration of 1.2 million cells/mL. Corresponds to the length of the green portion of the slab (live cells) over the total portion. * p≤0.05. (D) Dimensionless viability of MIN6 cells at a concentration of 10 million cells/mL. (error bar represents the standard error of the mean, N=3)

The dimensionless viability is the ratio of depth of living cells over the entire depth of the slab. The depth of living cells is determined by measuring the length from the top of the slab to where the living (green) cells abruptly changes to dead (red) cells as shown in **Figure 7A** and **Figure 7B**. The difference between hydrogels at high cell fractions is less noticeable (**Figure 7D**) compared to at a lower cell fraction (**Figure 7C**) where a significant difference was noticed among groups (p<0.05). As predicted by the trends in the effectiveness factor, changes in viability would become less noticeable at a higher Thiele modulus and therefore cell fraction by extension.

Table 3 presents the effectiveness factors and Thiele moduli for all hydrogels, calculated at the cell fractions used in the experiments shown in **Figure 7**. As expected, at a concentration of 10 million cells/mL, the differences in effectiveness factor between groups are minimal, due to the high Thiele modulus being much larger than the critical value. However, at a concentration of 1.2 million cells/mL, significant differences are observed: 2% Protanal reportedly achieves an effectiveness factor of 1, while 5% Manugel theoretically achieves only an effectiveness factor of 0.7 at the same cell fraction.

**Table 3.** Thiele modulus and effectiveness factor at 1.2 million cells/mL and 10 million cells/mL for MIN6.

| Hydrogel | 1.2 million MIN6 cells/mL |  | 10 million MIN6 cells/mL |  |
| --- | --- | --- | --- | --- |
| | Thiele Modulus ($\phi$) | Effectiveness factor ($\eta$) | Thiele Modulus ($\phi$) | Effectiveness factor ($\eta$) |
| 2% Manugel | 1.7 | 0.83 | 7.6 | 0.19 |
| 2% Protanal | 1.3 | 1 | 6.6 | 0.21 |
| 5% Manugel | 2.0 | 0.72 | 8.6 | 0.16 |

## 4. Discussion

In this work, we developed a simple platform to assess oxygen diffusion coefficient in hydrogels such as alginate. Oxygen diffusion coefficients determined using this setup were applied to predict the effect of beta cell seeding density on cell survival, which correlated with simplified unidimensional oxygen diffusion models.

First, the system was validated using water at 37°C as a reference, resulting in an experimental diffusion coefficient of 3.2×10^-5^ cm^2^s^-1^ ± 0.5×10^-5^ cm^2^s^-1^, which falls within the range of documented values ranging between 2.6×10^-5^ cm^2^s^-1^ [42] and 3.83×10^-5^ cm^2^s^-1^ [55]. Experimental estimates were obtained through transient counter-diffusion of oxygen after a step change from air to pure oxygen applied to the system. Experimental trends deviated from predicted curves for unidirectional diffusion-driven mass transport models, which is characteristic of larger Peclet number systems [52]. Accounting for advection could improve the accuracy of measured diffusion coefficients for fluids such as water. However, advection should be less problematic with hydrogels where fluid motion is hindered by the gel material. The geometry, gas flow, or temperature control strategies could potentially be refined to further reduce sources of error and advection effects.

The system was then applied to various formulations of alginate to determine their diffusion properties. Interestingly, the different formulations of alginate exhibited a much more significant impact on the measured diffusivity coefficient than the concentration of alginate itself. This observation suggests that polymer and gel properties such as the molecular weight distribution, block structure, and degree of crosslinking, can profoundly influence the diffusion characteristics. Since the gels were prepared through internal gelation, the formation of CO_2_ bubbles could also impact the gel rheological and gas diffusion properties. Despite the chemical composition and chain lengths of the different alginate types appearing nearly identical, subtle variations in the microstructure and crosslinking density may have led changes in polymer microstructure that impact oxygen diffusion properties. This finding emphasizes the need for experimental measurement of the diffusion coefficient, as theoretical predictions based solely on concentration or viscosity may overlook critical microstructural differences that affect transport properties. Such variations in the microstructure can lead to discrepancies between estimated and actual diffusion coefficients, which can be critical in applications where precise control over diffusion is required.

The effectiveness factor (η) can be calculated from the Thiele modulus and is a non-dimensional number between 0 and 1 that helps understand how much a system is limited by mass transfer, particularly in describing how well cells are being oxygenated. If oxygen is abundant and can travel freely into the cells, the effectiveness would be 1. Conversely, if oxygen cannot diffuse fast enough into the cells due to mass transfer limitations of the material, the effectiveness factor would be low (close to 0). The effect of diffusion on cell viability was quantified by live/dead staining and related to the Thiele modulus, which indicates the rate of cellular oxygen consumption and determines whether an encapsulation device/system can meet this demand or is limited by mass transfer and diffusion. At high cell fractions (10 million MIN6 cells/mL) where the Thiele modulus is large (φ>>φcrit), cells become hypoxic because oxygen cannot diffuse through the material fast enough to meet their demand, leading to necrosis and significantly reduced effectiveness (η<<1). In the live/dead experiment (**Figure 7C** and **Figure 7D**), the oxygen within the material is not replenishing fast enough and decreases rapidly as a function of depth resulting in a very sharp live/dead cut-off (η<<1). At such a high Thiele modulus like in **Figure 7D**, the difference between the effectiveness factors for different materials becomes smaller. At a lower cell fraction (1.2 million cells/mL), differences between the materials become more obvious showing a significant difference between 2% protanal and 5% Manugel (see **Figure 7B**).

The theoretical cell fractions in **Table 1** suggest that the 2% protanal sample should have had adequate oxygenation at 1.2 million cells/mL which was not the case. This suggests that the model needs refinement when compared with the results in **Figure 7**. Improved measure of the oxygen consumption rate or including Michaelis-Menten kinetics could overcome model limitations. Nevertheless, the results support the Thiele modulus hypothesis, where the difference in viability between samples is larger at low cell densities and much less obvious at high cell densities.

The data from **Figure 7D** suggests that in a system with a large Thiele modulus (φ>>φ_crit_), the oxygenation and cell viability are determined more by the cell fraction and oxygen consumption rate (cell type) than the choice of encapsulation material. At lower cell fractions where the effectiveness is closer to 1, the choice of material becomes more important.

Like MIN6 cells, islets are also regarded as a cell type with high oxidative capacity, natively they account for 10% of the blood flow in pancreas while only making up 1-2% of the cell mass [56]. At the cell fraction needed for a therapeutic implantable device (∼600,000 IEQ for a 60kg person) [57] in a small device (<10mL), we hypothesize the therapeutic implantable device encapsulation material choice will have little impact on cell viability which will be low in all conditions.

A microencapsulation device with a volume of approximately 375 mL is required to ensure the viability of a therapeutic dose of islets that are adequately oxygenated through diffusion (**Table 1**) (600,000 islets at ∼1600 islets/mL for 2% Protanal with η = 0.9). This size limitation can be addressed using alternative oxygenation strategies, such as advective transport through flow, oxygen-carrying fluids like blood, or incorporating oxygen-generating beads into the material [16,25]. In these cases, the choice of encapsulation material and its corresponding oxygen diffusion coefficient becomes increasingly critical, as the system may behave similarly to the scenario observed in **Figure 7C** with MIN6 cells at lower cell densities. A more diffusive material could enable a smaller device with equivalent performance and the same therapeutic insulin dose. In islet transplantation, it is common to use an excess of islets to compensate for early cell death due to poor oxygenation. If cells are adequately oxygenated (η≥0.9), the overall cell fraction and device size could be reduced, as compensation for early cell death would be unnecessary. Minimizing device size and cell fraction through improved oxygen diffusion could increase the availability of donated islets for transplantation, thereby expanding access to this treatment.

## 5. Conclusion

A simple method to measure the oxygen diffusion coefficient was developed for liquids and hydrogels commonly used for cellular encapsulation. The system was validated and measured the diffusion coefficient of water as 3.2×10^-5^ ± 0.5×10^-5^ cm^2^s^-1^ at 37°C which is within the range of what is currently documented in literature. The system measured the diffusion coefficient of three different formulations of hydrogels (2% Manugel, 2% Protanal, and 5% Manugel). Our study demonstrates a clear correlation between the loss modulus of hydrogels and their effective diffusion coefficients, with lower loss modulus values corresponding to higher diffusion rates. These findings emphasize the critical role of rheological properties in influencing diffusion behavior in hydrogel systems. Moreover, measuring the oxygen diffusivity experimentally allows a more predictive approximation of Thiele and effectiveness factors which are useful dimensionless numbers for encapsulation device sizing and predicting trends is cellular viability.

## Supporting information

STL files

COMSOL files

## Acknowledgements

The authors gratefully acknowledge the funding that made this work possible, including the Stem Cell Network Fueling Biotechnology grant in collaboration with Aspect Biosystems, led by Timothy Kieffer at UBC. The work was also supported by funding from JDRF, Canadian Institutes of Health Research, Diabetes Canada, NSERC Discovery, and the Canada Research Chairs program (C.H.). We also acknowledge the support provided by the following networks: the Quebec Cell, Tissue and Gene Therapy Network – ThéCell, the Cardiometabolic Health, Diabetes and Obesity – CMDO Research Network (thematic networks supported by the Fonds de recherche du Québec – Santé), PROTEO (The Quebec Network for Research on Protein Function), and CQMF/QCAM (Quebec Centre for Advanced Materials). Additionally, we would like to acknowledge the funding from the Vadasz Scholar Award from McGill University (K.C.) and the Fonds de recherche du Québec – Nature et Technologies (FRQNT). We thank Florent Lemaire and Marco Pineda for their input in experimental design and analysis. We also gratefully acknowledge Wenche Iren Strand at Norwegian Biopolymer Laboratory (NOBIPOL), Department of Biotechnology and Food Science, Norwegian University of Science and Technology for the analysis of the alginates.

---

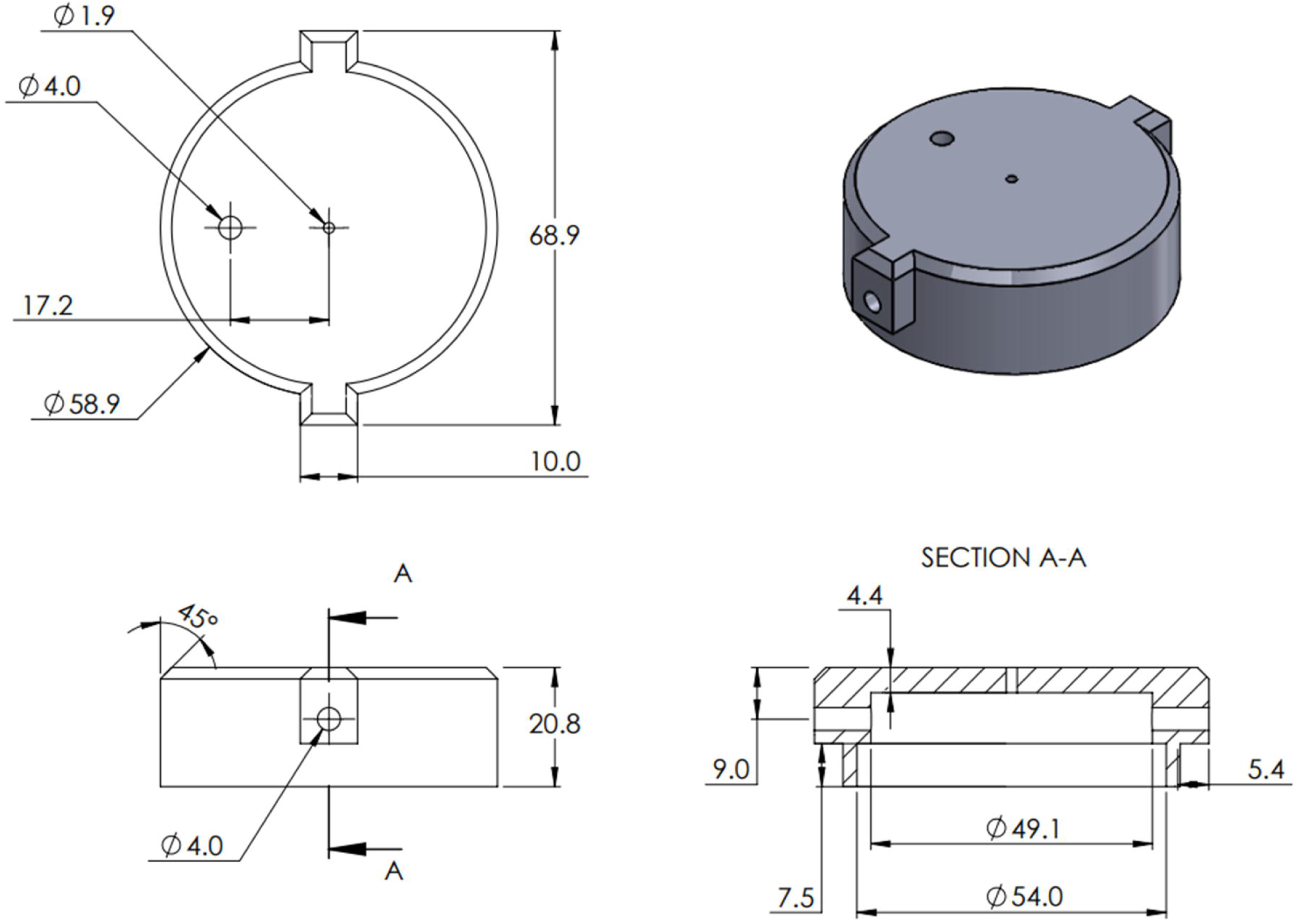

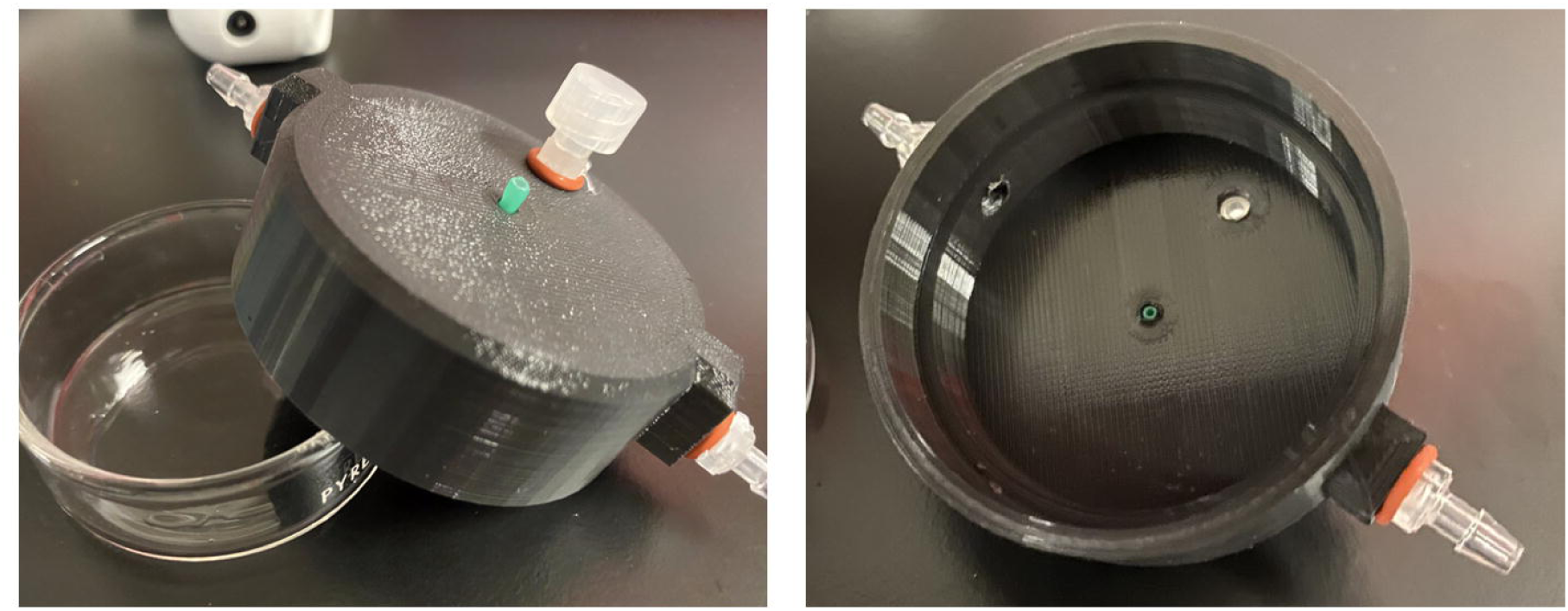

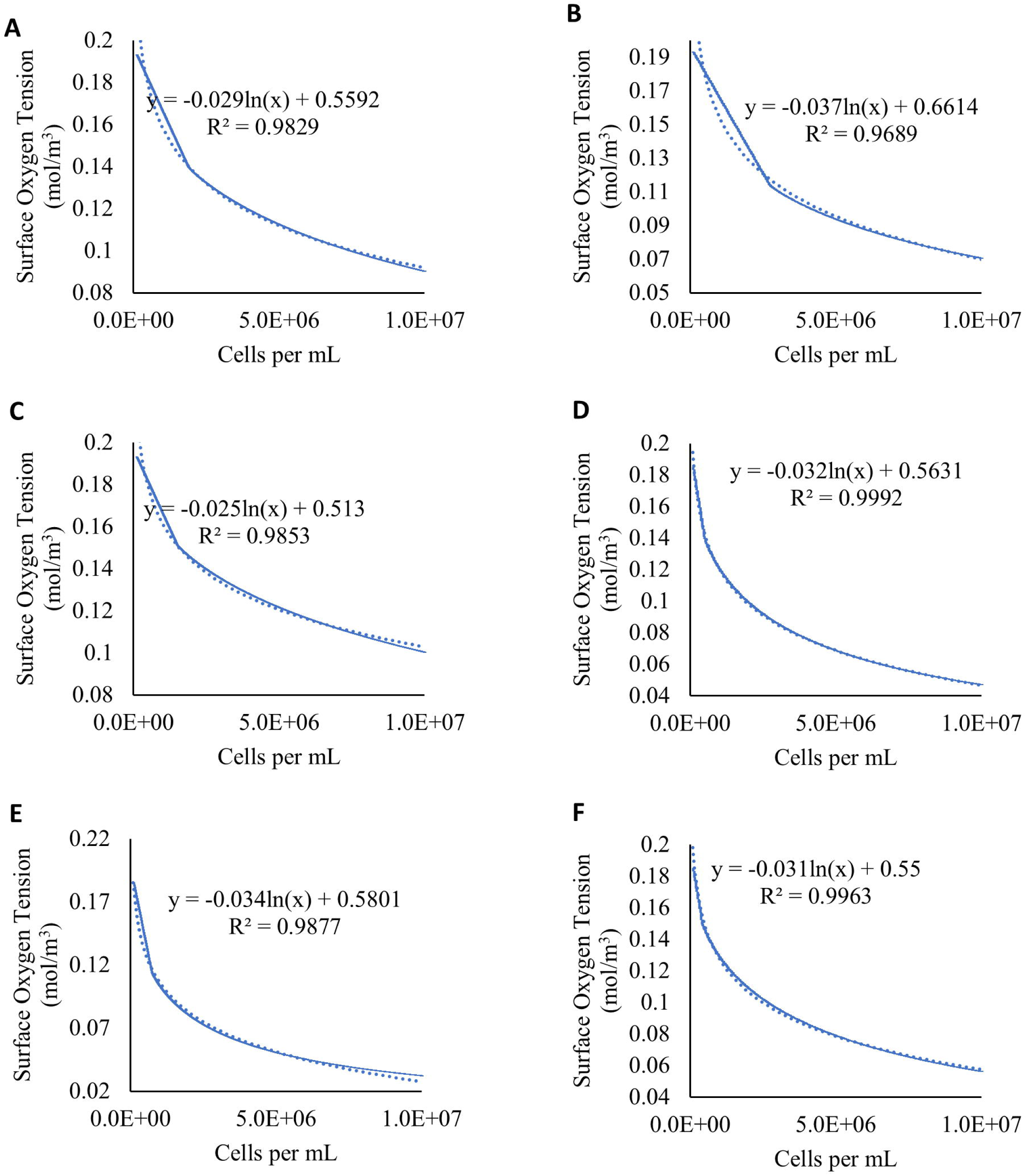

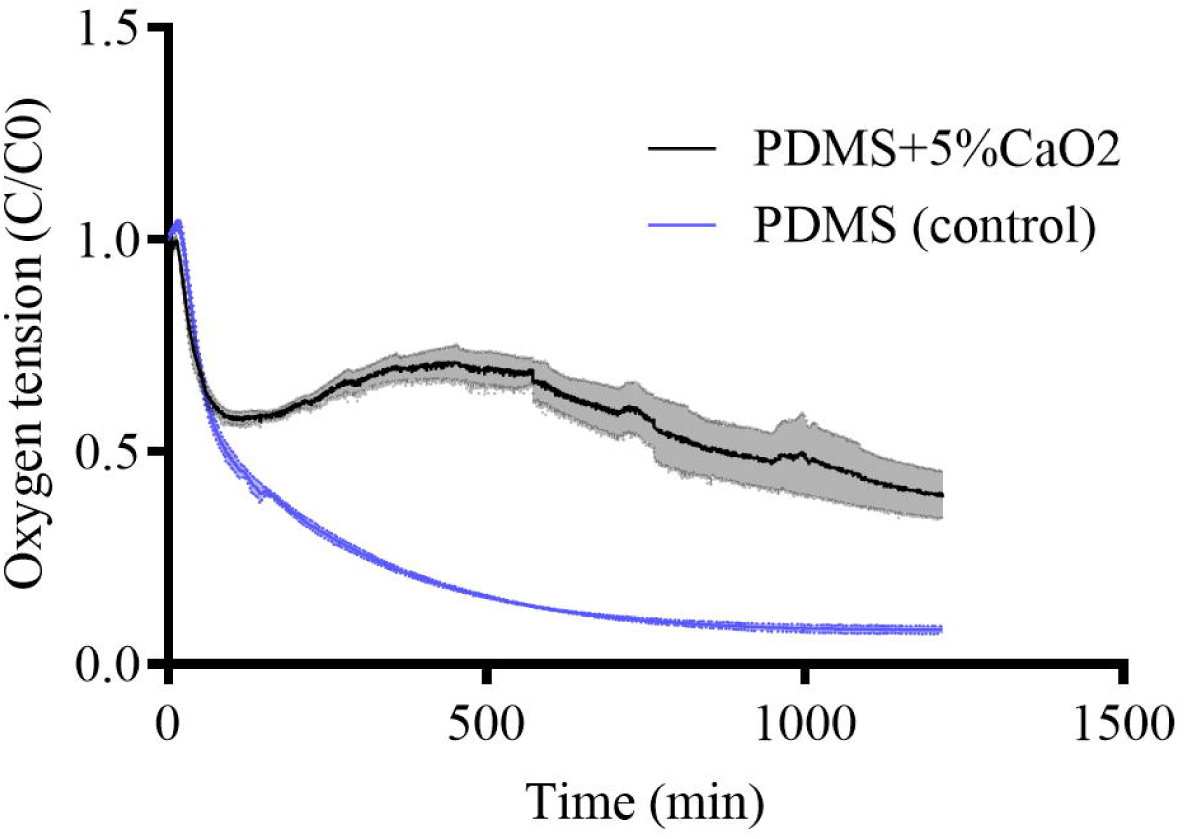

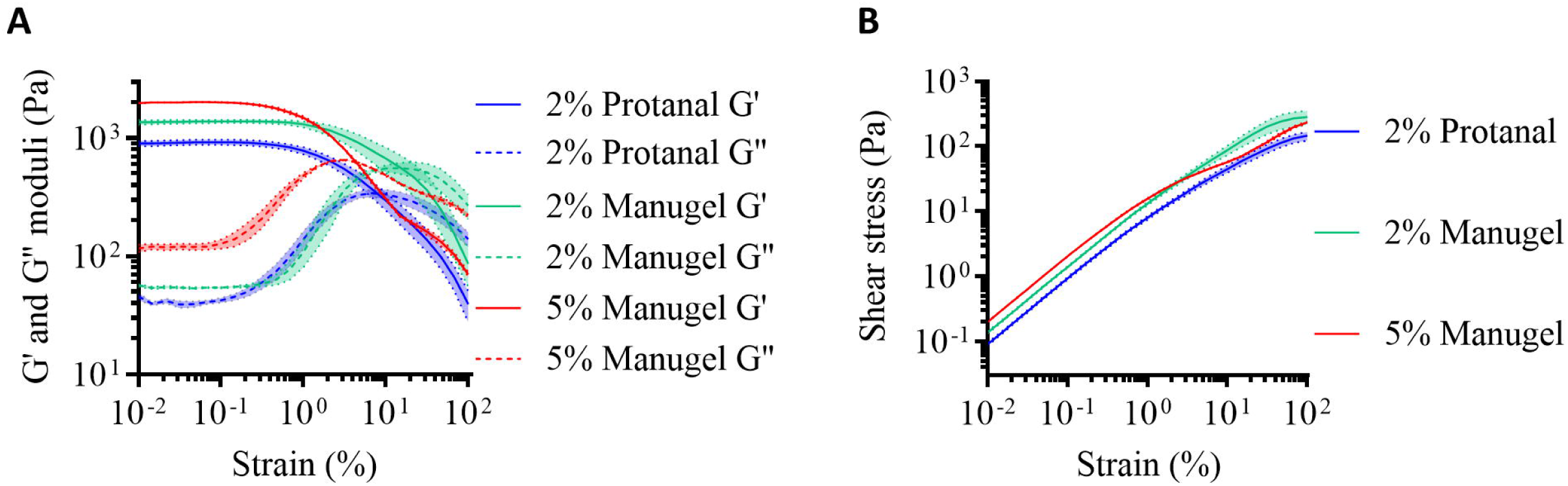

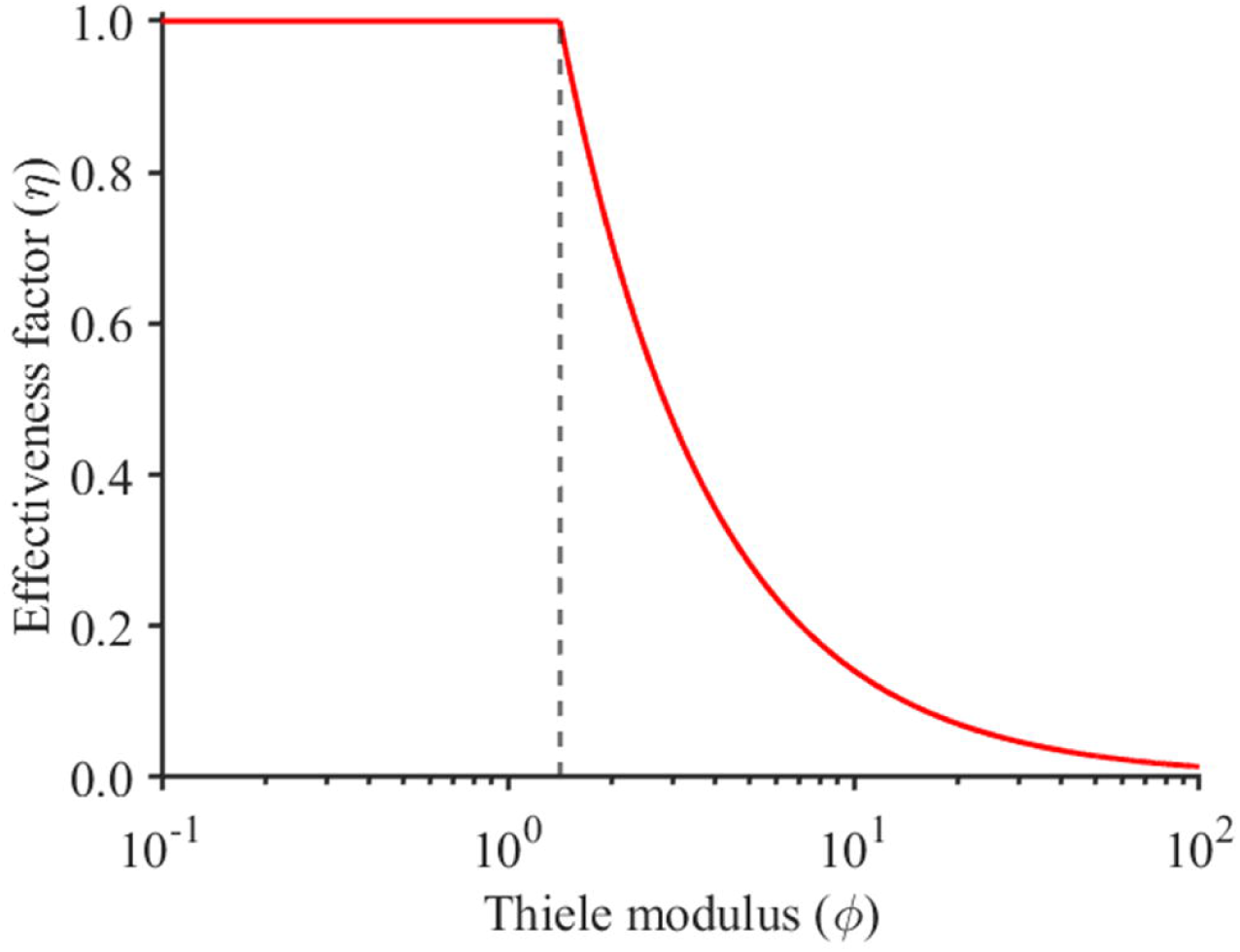

## References

[1] Thomas M.S. Chang, Semipermeable Microcapsules, Science (80-. ). 146 (1964) 1–2. 10.1126/science.146.3643.524.

[2] D.I.C. Biotechnol, O. Smidsrd, Alginate as immobilization matrix for cells, Trends Biotechnol. 8 (1990). 10.1016/0167-7799(90)90139-o.

[3] C.A. Hoesli, M. Luu, J.M. Piret, A novel alginate hollow fiber bioreactor process for cellular therapy applications, Biotechnol. Prog. 25 (2009) 1740–1751. 10.1002/btpr.260.

[4] L.K. Narayanan, P. Huebner, M.B. Fisher, J.T. Spang, B. Starly, R.A. Shirwaiker, 3D- Bioprinting of Polylactic Acid (PLA) Nanofiber–Alginate Hydrogel Bioink Containing Human Adipose-Derived Stem Cells, ACS Biomater. Sci. Eng. 2 (2016) 1732–1742. 10.1021/acsbiomaterials.6b00196.

[5] S.D. Eswaramoorthy, N. Dhiman, A. Joshi, S.N. Rath, 3D Bioprinting of Mesenchymal Stem Cells and Endothelial Cells in an Alginate-Gelatin-Based Bioink, J. 3D Print. Med. 5 (2021) 23–36. 10.2217/3dp-2020-0026.

[6] J. Jia, D.J. Richards, S. Pollard, Y. Tan, J. Rodriguez, R.P. Visconti, T.C. Trusk, M.J. Yost, H. Yao, R.R. Markwald, Y. Mei, Acta Biomaterialia Engineering alginate as bioink for bioprinting, Acta Biomater. 10 (2014) 4323–4331. 10.1016/j.actbio.2014.06.034.

[7] K. Ziv, H. Nuhn, Y. Ben-haim, L.S. Sasportas, P.J. Kempen, T.P. Niedringhaus, M. Hrynyk, R. Sinclair, A.E. Barron, S.S. Gambhir, Biomaterials A tunable silk e alginate hydrogel scaffold for stem cell culture and transplantation, Biomaterials. 35 (2014) 3736– 3743. 10.1016/j.biomaterials.2014.01.029.

[8] E. Trouche, S.G. Fullana, C. Mias, C. Ceccaldi, F. Tortosa, M.H. Seguelas, D. Calise, A. Parini, D. Cussac, B. Sallerin, Evaluation of Alginate Microspheres for Mesenchymal Stem Cell Engraftment on Solid Organ, Cell Transplant. 19 (2010) 1623–1633. 10.3727/096368910X514297.

[9] J. Yu, K.T. Du, Q. Fang, Y. Gu, S.S. Mihardja, R.E. Sievers, J.C. Wu, R.J. Lee, Biomaterials The use of human mesenchymal stem cells encapsulated in RGD modi fi ed alginate microspheres in the repair of myocardial infarction in the rat, Biomaterials. 31 (2010) 7012–7020. 10.1016/j.biomaterials.2010.05.078.

[10] D.A. Alagpulinsa, J.J.L. Cao, R.K. Driscoll, R.F. Sîrbulescu, M.F.E. Penson, M. Sremac, E.N. Engquist, T.A. Brauns, J.F. Markmann, D.A. Melton, M.C. Poznansky, Alginate - microencapsulation of human stem cell – derived β cells with CXCL12 prolongs their survival and function in immunocompetent mice without systemic immunosuppression, Am. J. Transplant. 19 (2019) 1930–1940. 10.1111/ajt.15308.

[11] S. Swioklo, P. Ding, A.W. Pacek, C.J. Connon, Process parameters for the high-scale production of alginate-encapsulated stem cells for storage and distribution throughout the cell therapy supply chain, Process Biochem. 59 (2017) 289–296. 10.1016/j.procbio.2016.06.005.

[12] M. Neumann, T. Arnould, B.-L. Su, Encapsulation of stem-cell derived β-cells: A promising approach for the treatment for type 1 diabetes mellitus, J. Colloid Interface Sci. 636 (2023) 90–102. 10.1016/j.jcis.2022.12.123.

[13] M. Farina, J.F. Alexander, U. Thekkedath, M. Ferrari, A. Grattoni, Cell encapsulation: Overcoming barriers in cell transplantation in diabetes and beyond, Adv. Drug Deliv. Rev. 139 (2019) 92–115. 10.1016/j.addr.2018.04.018.

[14] K. Shah, Encapsulated stem cells for cancer therapy, Biomatter. 3 (2013). 10.4161/biom.24278.

[15] C.K. Colton, Oxygen supply to encapsulated therapeutic cells, Adv. Drug Deliv. Rev. 67– 68 (2014) 93–110. 10.1016/j.addr.2014.02.007.

[16] and H.C. Fernandez SA, Champion KS, Leask RL, Engineering Vascularized Islet Macroencapsulation Devices: An in vitro Platform to Study Oxygen Transport in Perfused Immobilized Pancreatic Beta Cell Cultures, Front. Bioeng. Biotechnol. 10 (2022) 884071. 10.3389/fbioe.2022.884071.

[17] H. Komatsu, F. Kandeel, Y. Mullen, Impact of Oxygen on Pancreatic Islet Survival, Pancreas. 47 (2018) 533–543. 10.1097/MPA.0000000000001050.

[18] D.T. Bowers, W. Song, L.-H. Wang, M. Ma, Engineering the vasculature for islet transplantation, Acta Biomater. 95 (2019) 131–151. 10.1016/j.actbio.2019.05.051.

[19] T. Linn, J. Schmitz, I. Hauck-Schmalenberger, Y. Lai, R.G. Bretzel, H. Brandhorst, D. Brandhorst, Ischaemia is linked to inflammation and induction of angiogenesis in pancreatic islets, Clin. Exp. Immunol. 144 (2006) 179–187. 10.1111/j.1365-2249.2006.03066.x.

[20] B.N. Moeun, S. Da Ling, M. Gasparrini, A.K. Rutman, S. Negi, S. Paraskevas, C.A. Hoesli, Islet Encapsulation: A Long-Term Treatment for Type 1 Diabetes, in: Ref. Modul. Biomed. Sci., Elsevier, 2019. 10.1016/B978-0-12-801238-3.11135-3.

[21] and C.A.H. Fernandez, Stephanie, Marc-André Dussault, André Bégin-Drolet, Jean Ruel, R. Leask, Towards the fabrication of a 3D printed vascularized islet transplantation device for the treatment of type 1 diabetes, Front. Bioeng. Biotechnol. 4 (2016). 10.3389/conf.FBIOE.2016.01.01021.

[22] P. Buchwald, A local glucose-and oxygen concentration-based insulin secretion model for pancreatic islets, Theor. Biol. Med. Model. 8 (2011) 20. 10.1186/1742-4682-8-20.

[23] A.S. Johnson, R.J. Fisher, G.C. Weir, C.K. Colton, Oxygen consumption and diffusion in assemblages of respiring spheres: Performance enhancement of a bioartificial pancreas, Chem. Eng. Sci. 64 (2009) 4470–4487. 10.1016/j.ces.2009.06.028.

[24] R. Cao, E. Avgoustiniatos, K. Papas, P. de Vos, J.R.T. Lakey, Mathematical predictions of oxygen availability in micro- and macro-encapsulated human and porcine pancreatic islets, J. Biomed. Mater. Res. Part B Appl. Biomater. 108 (2020) 343–352. 10.1002/jbm.b.34393.

[25] J.-P. Liang, R.P. Accolla, M. Soundirarajan, A. Emerson, M.M. Coronel, C.L. Stabler, Engineering a macroporous oxygen-generating scaffold for enhancing islet cell transplantation within an extrahepatic site, Acta Biomater. 130 (2021) 268–280. 10.1016/j.actbio.2021.05.028.

[26] E.S. Avgoustiniatos, C.K. Colton, M. Of, C. Viability, Effect of External Oxygen Mass Transfer Resistances on Viability of Immunoisolated Tissuea, (n.d.).

[27] A. Al-Ani, D. Toms, D. Kondro, J. Thundathil, Y. Yu, M. Ungrin, Oxygenation in cell culture: Critical parameters for reproducibility are routinely not reported, PLoS One. 13 (2018) e0204269. 10.1371/journal.pone.0204269.

[28] Ü. Mehmetoglu, S. Ateş, R. Berber, Oxygen Diffusivity in Calcium Alginate Gel Beads Containing Gluconobacter Suboxydans, Artif. Cells, Blood Substitutes, Biotechnol. 24 (1996) 91–106. 10.3109/10731199609118877.

[29] D. Cristea, S. Krishtul, P. Kuppusamy, L. Baruch, M. Machluf, A. Blank, New approach to measuring oxygen diffusion and consumption in encapsulated living cells, based on electron spin resonance microscopy, Acta Biomater. 101 (2020) 384–394. 10.1016/j.actbio.2019.10.032.

[30] M. Kotecha, L. Wang, S. Hameed, N. Viswakarma, M. Ma, C. Stabler, C.A. Hoesli, B. Epel, In vitro oxygen imaging of acellular and cell - loaded beta cell replacement devices, Sci. Rep. (2023) 1–17. 10.1038/s41598-023-42099-w.

[31] L. Figueiredo, R. Pace, C. D’Arros, G. Réthoré, J. Guicheux, C. Le Visage, P. Weiss, Assessing glucose and oxygen diffusion in hydrogels for the rational design of 3D stem cell scaffolds in regenerative medicine, J. Tissue Eng. Regen. Med. 12 (2018) 1238–1246. 10.1002/term.2656.

[32] K. Bhunia, S.S. Sablani, J. Tang, B. Rasco, Non-invasive measurement of oxygen diffusion in model foods, Food Res. Int. 89 (2016) 161–168. 10.1016/j.foodres.2016.07.015.

[33] Y. Rharbi, A. Yekta, M.A. Winnik, A Method for Measuring Oxygen Diffusion and Oxygen Permeation in Polymer Films Based on Fluorescence Quenching, Anal. Chem. 71 (1999) 5045–5053. 10.1021/ac990193c.

[34] T. Fiedler, I. V. Belova, G.E. Murch, G. Poologasundarampillai, J.R. Jones, J.A. Roether, A.R. Boccaccini, A comparative study of oxygen diffusion in tissue engineering scaffolds, J. Mater. Sci. Mater. Med. 25 (2014) 2573–2578. 10.1007/s10856-014-5264-7.

[35] A.E. AL-Muftah, I.M. Abu-Reesh, Effects of internal mass transfer and product inhibition on a simulated immobilized enzyme-catalyzed reactor for lactose hydrolysis, Biochem. Eng. J. 23 (2005) 139–153. 10.1016/j.bej.2004.10.010.

[36] E.S. Avgoustiniatos, C.K. Colton, Effect of External Oxygen Mass Transfer Resistances on Viability of Immunoisolated Tissue a, Ann. N. Y. Acad. Sci. 831 (1997) 145–166. 10.1111/j.1749-6632.1997.tb52192.x.

[37] R. Olsson, P.-O. Carlsson, A Low-Oxygenated Subpopulation of Pancreatic Islets Constitutes a Functional Reserve of Endocrine Cells, Diabetes. 60 (2011) 2068–2075. 10.2337/db09-0877.

[38] J.R.H. Ross, Mass and Heat Transfer Limitations and Other Aspects of the Use of Large-Scale Catalytic Reactors, in: Contemp. Catal., Elsevier, 2019: pp. 187–213. 10.1016/B978-0-444-63474-0.00008-4.

[39] F.M. Pereira, S.C. Oliveira, Occurrence of dead core in catalytic particles containing immobilized enzymes: analysis for the Michaelis–Menten kinetics and assessment of numerical methods, Bioprocess Biosyst. Eng. 39 (2016) 1717–1727. 10.1007/s00449-016-1647-0.

[40] J. Miyazaki, K. Araki, E. Yamato, H. Ikegami, K. Yamamura, Establishment of a Pancreatic /? Cell Line That Retains Glucose-Inducible Insulin Secretion : Special Reference to Expression of Glucose Transporter Isoforms *, 127 (1990).

41. [41] X. Liu, stage_file_maker, (2023). https://www.mathworks.com/matlabcentral/fileexchange/68849-stage_file_maker.

[42] S. Preibisch, S. Saalfeld, P. Tomancak, Globally optimal stitching of tiled 3D microscopic image acquisitions, Bioinformatics. 25 (2009) 1463–1465. 10.1093/bioinformatics/btp184.

[43] H. Grasdalen, B. Larsen, O. Smidsrød, A p.m.r. study of the composition and sequence of uronate residues in alginates, Carbohydr. Res. 68 (1979) 23–31. 10.1016/S0008-6215(00)84051-3.

[44] H. Ertesvåg, G. Skjåk-Bræk, Modification of Alginate Using Mannuronan C-5- Epimerases, in: Carbohydr. Biotechnol. Protoc., 1999: pp. 71–78. 10.1007/978-1-59259-261-6_6.

[45] H. Grasdalen, High-field, 1H-n.m.r. spectroscopy of alginate: sequential structure and linkage conformations, Carbohydr. Res. 118 (1983) 255–260. 10.1016/0008-6215(83)88053-7.

[46] I.M.N. Vold, K.A. Kristiansen, B.E. Christensen, A Study of the Chain Stiffness and Extension of Alginates, in Vitro Epimerized Alginates, and Periodate-Oxidized Alginates Using Size-Exclusion Chromatography Combined with Light Scattering and Viscosity Detectors, Biomacromolecules. 7 (2006) 2136–2146. 10.1021/bm060099n.

[47] C.R. Wilke, P. Chang, Correlation of diffusion coefficients in dilute solutions, AIChE J. 1 (1955) 264–270. 10.1002/aic.690010222.

[48] F. Noël, B. Mauroy, Interplay Between Optimal Ventilation and Gas Transport in a Model of the Human Lung, Front. Physiol. 10 (2019). 10.3389/fphys.2019.00488.

[49] P. Buchwald, FEM-based oxygen consumption and cell viability models for avascular pancreatic islets, Theor. Biol. Med. Model. 6 (2009) 5. 10.1186/1742-4682-6-5.

[50] W. Xing, M. Yin, Q. Lv, Y. Hu, C. Liu, J. Zhang, Oxygen Solubility, Diffusion Coefficient, and Solution Viscosity, in: Rotating Electrode Methods Oxyg. Reduct. Electrocatal., Elsevier, 2014: pp. 1–31. 10.1016/B978-0-444-63278-4.00001-X.

[51] S. Arora, F. Potůček, Modelling of displacement washing of packed bed of fibers, Brazilian J. Chem. Eng. 26 (2009) 385–393. 10.1590/S0104-66322009000200016.

[52] I.A. Ganaie, S. Arora, V.K. Kukreja, Modelling and Simulation of a Packed Bed of Pulp Fibers Using Mixed Collocation Method, Int. J. Differ. Equations. 2013 (2013) 1–7. 10.1155/2013/875298.

[53] A. Najdahmadi, J.R.T. Lakey, E. Botvinick, Structural Characteristics and Diffusion Coefficient of Alginate Hydrogels Used for Cell Based Drug Delivery, MRS Adv. 3 (2018) 2399–2408. 10.1557/adv.2018.455.

[54] B. Bellich, M. Borgogna, M. Cok, A. Cesàro, Release Properties of Hydrogels: Water Evaporation from Alginate Gel Beads, Food Biophys. 6 (2011) 259–266. 10.1007/s11483-011-9206-3.

[55] C. Androjna, J.E. Gatica, J.M. Belovich, K.A. Derwin, Oxygen Diffusion through Natural Extracellular Matrices: Implications for Estimating “Critical Thickness” Values in Tendon Tissue Engineering, Tissue Eng. Part A. 14 (2008) 559–569. 10.1089/tea.2006.0361.

[56] H. Peiris, C.S. Bonder, P.T.H. Coates, D.J. Keating, C.F. Jessup, The β-Cell/EC Axis: How Do Islet Cells Talk to Each Other?, Diabetes. 63 (2014) 3–11. 10.2337/db13-0617.

[57] S. Lablanche, S. Borot, A. Wojtusciszyn, K. Skaare, A. Penfornis, P. Malvezzi, L. Badet, C. Thivolet, E. Morelon, F. Buron, E. Renard, I. Tauveron, O. Villard, M. Munch, S. Sommacal, L. Clouaire, M. Jacquet, L. Gonsaud, C. Camillo-Brault, C. Colin, J.-L. Bosson, D. Bosco, T. Berney, L. Kessler, P.-Y. Benhamou, Ten-year outcomes of islet transplantation in patients with type 1 diabetes: Data from the Swiss-French GRAGIL network, Am. J. Transplant. 21 (2021) 3725–3733. 10.1111/ajt.16637.

