## Supplementary material for "An accessible platform to quantify oxygen diffusion in cell-laden hydrogels and its application to alginate-immobilized pancreatic beta cells": Text

Date: August 19, 2024

### 1 Technical Diagram of gas flow cap used in diffusion sensing setup

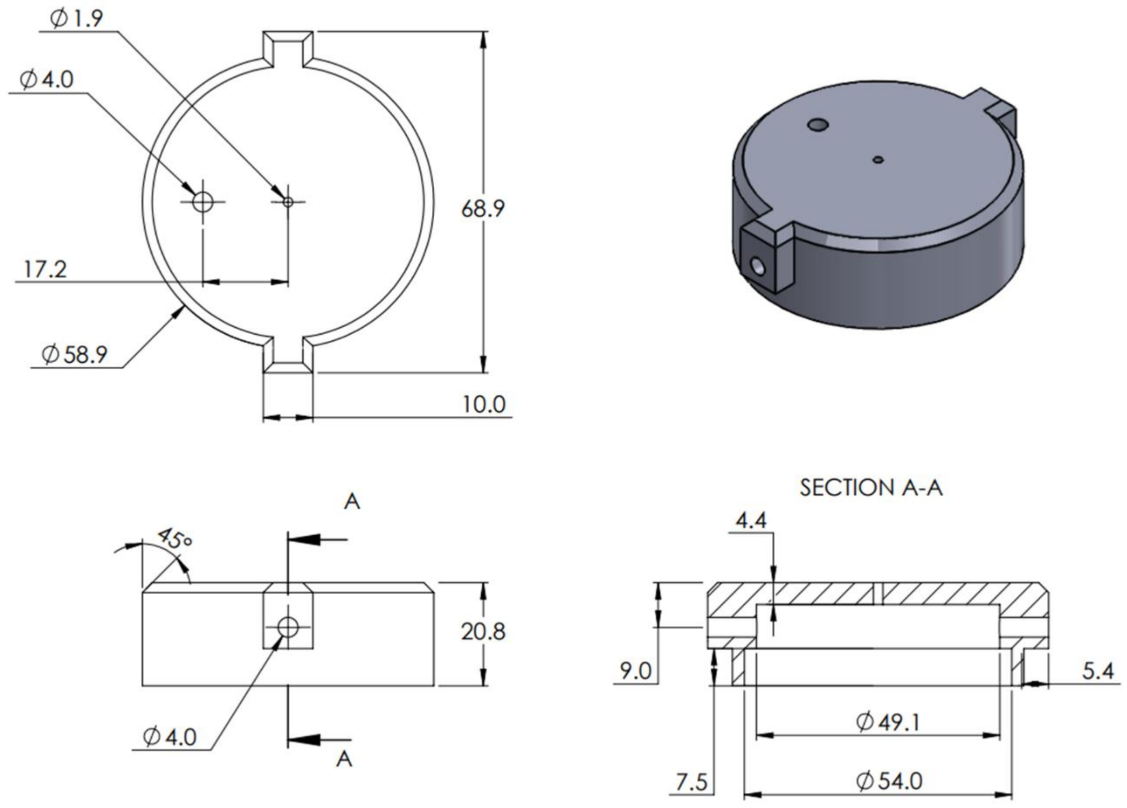


Figure S1. Technical drawing of gas flow cap used in diffusion sensing setup

Units: millimeters (mm)

Note: The STL file will be provided with publication and/or upon request.

Barbed tube hose fittings are attached to both ends; they can be manually forced through when hose fitting has an M4 thread. A Luer lock to hose thread can be attached to the top in the same manner. Between all fittings, small 4 mm O-rings were used to create a better seal. The Luer lock connection is sealed with a plastic male Luer lock fixture. Assembly pictures are shown below:

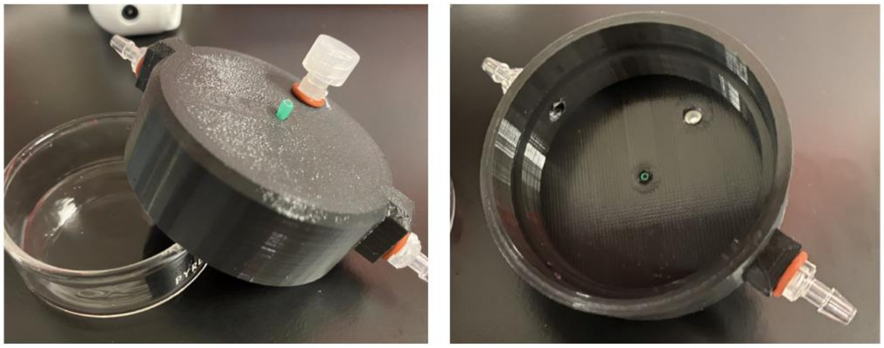

Figure S2. gas flow cap used in diffusion sensing setup

### 2 Relationship between surface concentration and cell fraction used for calculating the Thiele modulus

The oxygen tension at the surface of the alginate which is below a small layer of media could vary from the oxygen tension at the surface of the media.

A 1-D model following the geometry in Fig 3-1D was developed. In the media portion (total length: 1.5 mm), the diffusivity was assumed to be that of water at 37°C ($3.1\times10^{-5}$ $cm^2s^{-1}$). In the hydrogel, the effective diffusivity ($D_{eff}$) was calculated using a weighted average of the cell fraction (the $X$ variable), the oxygen diffusivity of tissue ($D_{tissue}$), and the experimental oxygen diffusivity of the hydrogel in question (2% Manugel, 2% Protanal, or 5% Manugel) referred to as $D_{alg}$. The oxygen diffusivity of tissue was taken as $1.24\times10^{-5}$ $cm^2s^{-1}$.

$$D_{eff} = (1 - X)D_{alg} + (X)D_{tissue}$$

Choosing either the OCR for islets or MIN6, a relationship between the cell fraction and surface oxygen tension can be developed for a known geometry.

Note that 0.44 mmHg corresponds to about $6\times10^{-4}$ $mol\cdot m^{-3}$, the Michaelis-Menten coefficient of primary human islets.

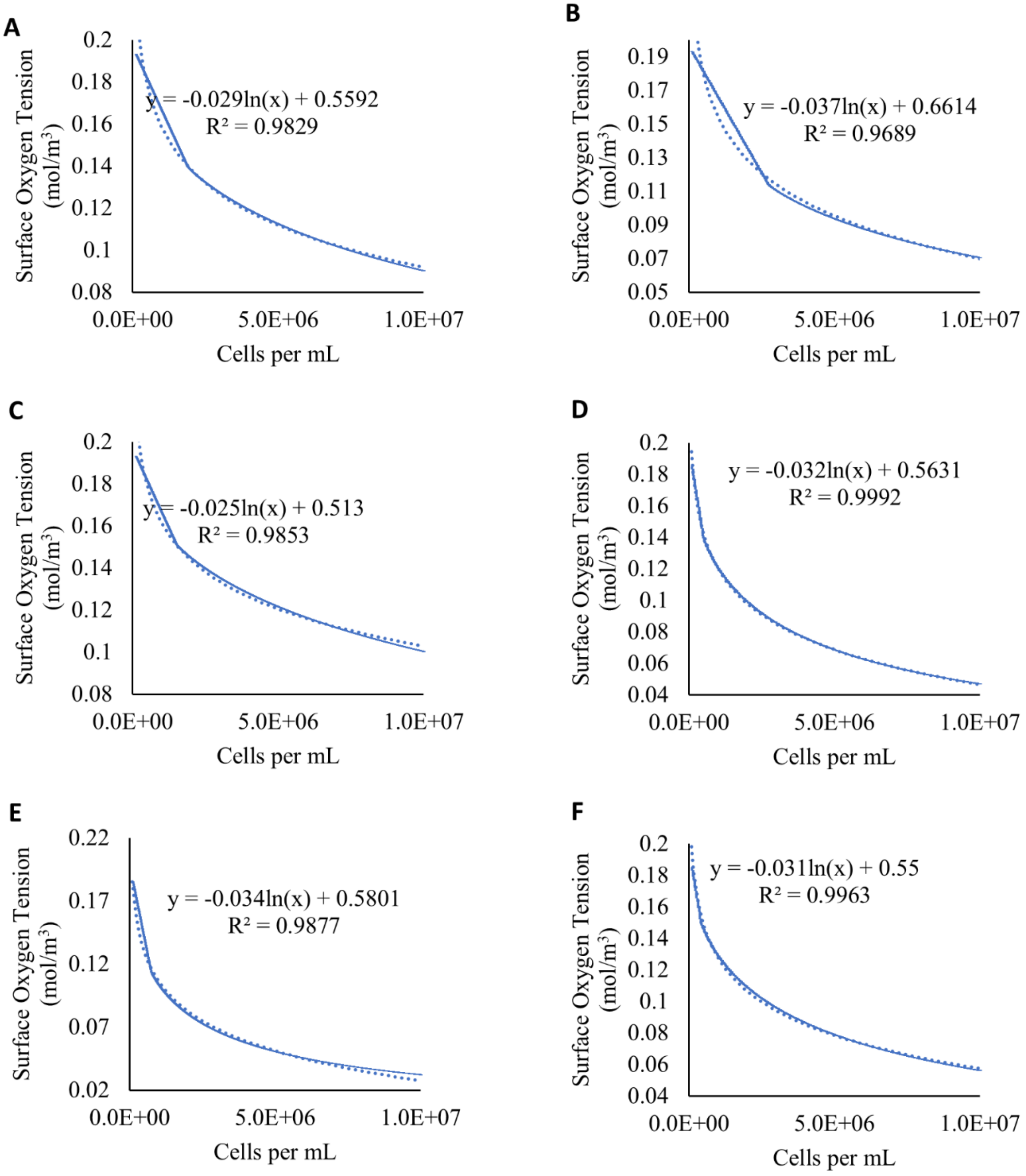


Figure S3. Surface oxygen tension of (A) 2% Manugel (B) 2% Protanal (C) 5% Manugel using human islets (OCR = 0.034 molm$^{-3}$s$^{-1}$), and (D) 2% Manugel (E) 2% Protanal (F) 5% Manugel using human islets (OCR = 0.129 molm$^{-3}$s$^{-1}$)

### 3 Monitoring oxygen release during oxygen purging process

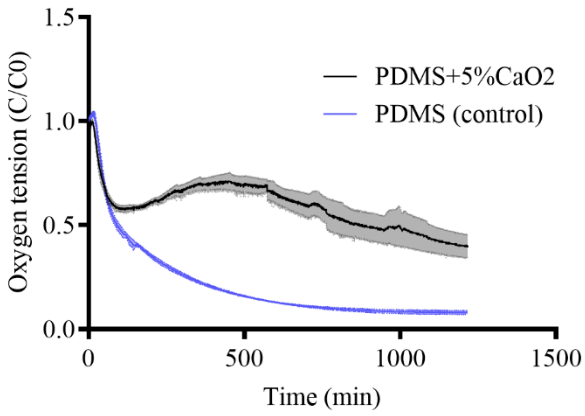


Figure S4. Experimental data of non-dimensional oxygen concentration in air-saturated water on PDMS + 5% $CaO_2$ (gray) and polydimethylsiloxane (PDMS, blue).

### 4 Amplitude sweep experiment in rheology of hydrogels at room temperature

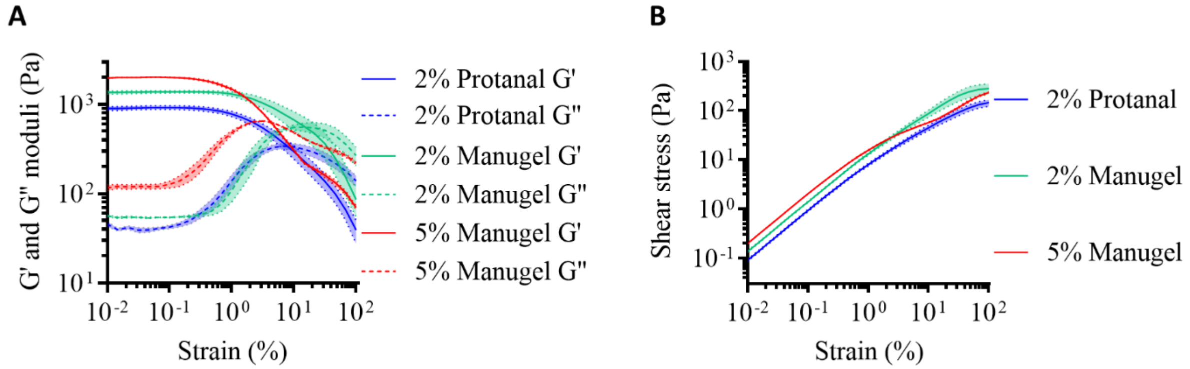


Figure S5. Rheological characterization of hydrogels. (A) Experimental storage and loss moduli as a function of strain (0.01 to 100 %) (B) Experimental shear stress versus strain (0.01 to 100 %) for 5% w/v Manugel alginate (red zone), 2% w/v Manugel alginate (green zone) and 2% w/v Protanal alginate (blue zone). (Oscillation frequency =10 1/s)

### 5 Effectiveness factor for 0th order reaction

Using the equation for the Thiele modulus and effectiveness factor, a theoretical estimation of the maximum cell fraction was determined for different hydrogels and cell types (MIN6 cells vs. islets). An effectiveness factor of 0.9 was chosen as the maximum allowable cell fraction since it theoretically corresponds to a fairly good level of oxygenation where the system is not limited by oxygen diffusion.

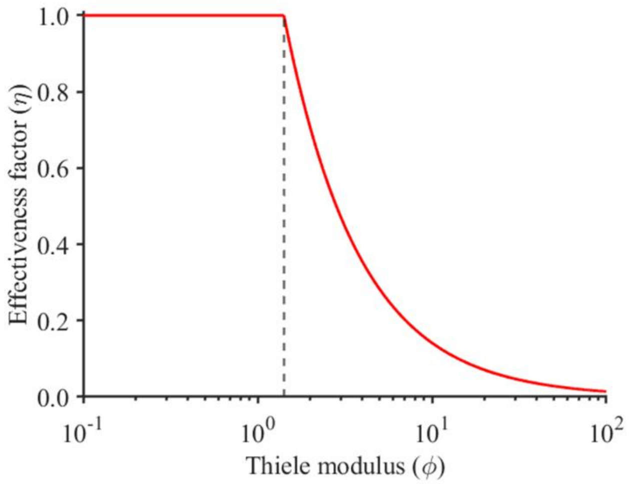


Figure S6. the effectiveness factor vs. Thiele modulus plot for a 0th order reaction such as the oxygen consumption of cells when $C_s >> K_m$. Note that the dashed line corresponds to the critical Thiele modulus ($\varphi_{crit}$) where the effectiveness begins decreasing from 1.
