## Supplementary material for "An accessible platform to quantify oxygen diffusion in cell-laden hydrogels and its application to alginate-immobilized pancreatic beta cells": COMSOL files: 20220912_modelling_figure extension.pptx

### Slide 1
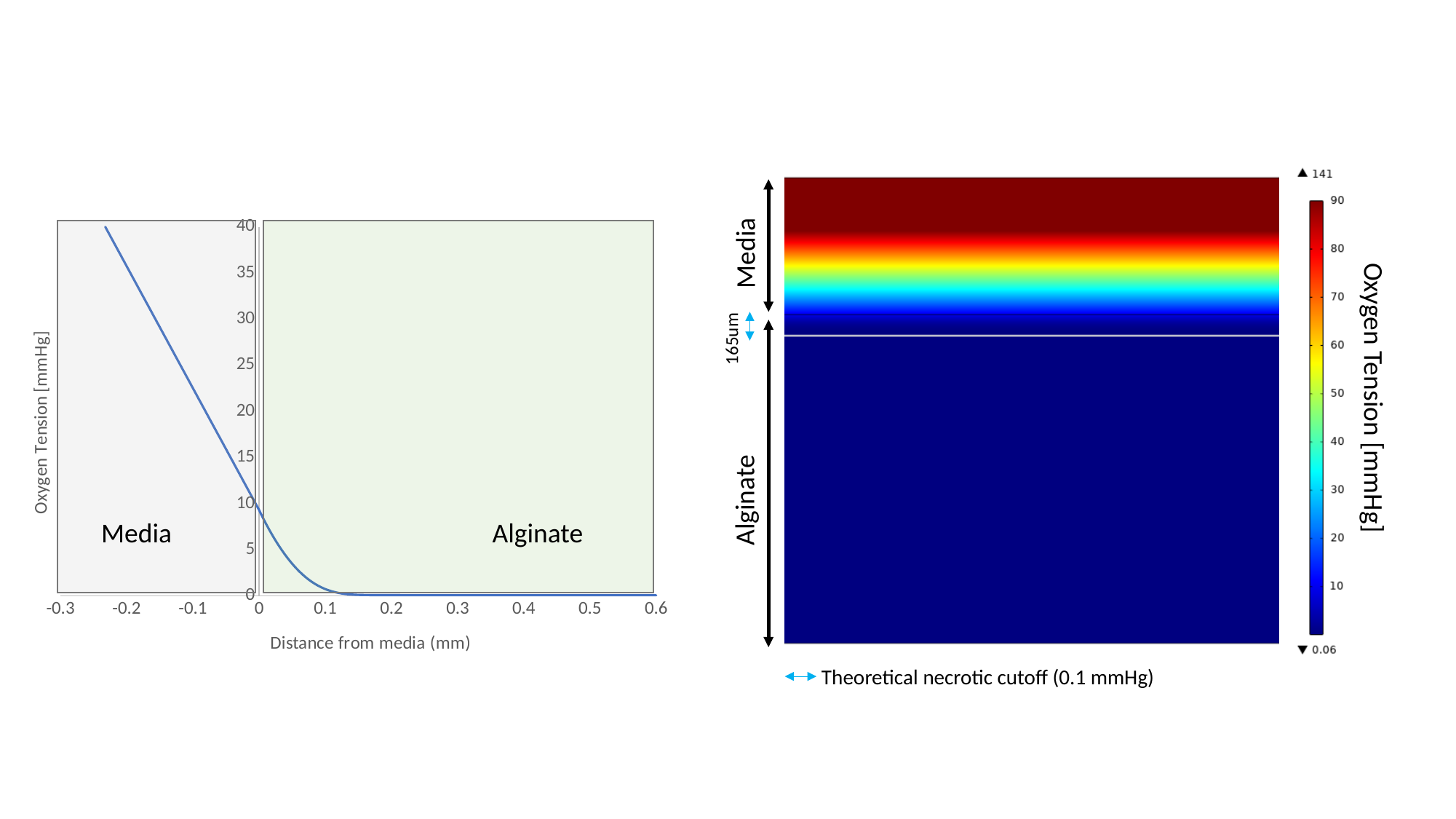

#### Chart
| Category | |
|---|---|Media
165um
Oxygen Tension [mmHg]
Alginate
Media
Alginate
Theoretical necrotic cutoff (0.1 mmHg)
