## Supplementary figures and images for "An accessible platform to quantify oxygen diffusion in cell-laden hydrogels and its application to alginate-immobilized pancreatic beta cells"

### Peclet_number.png

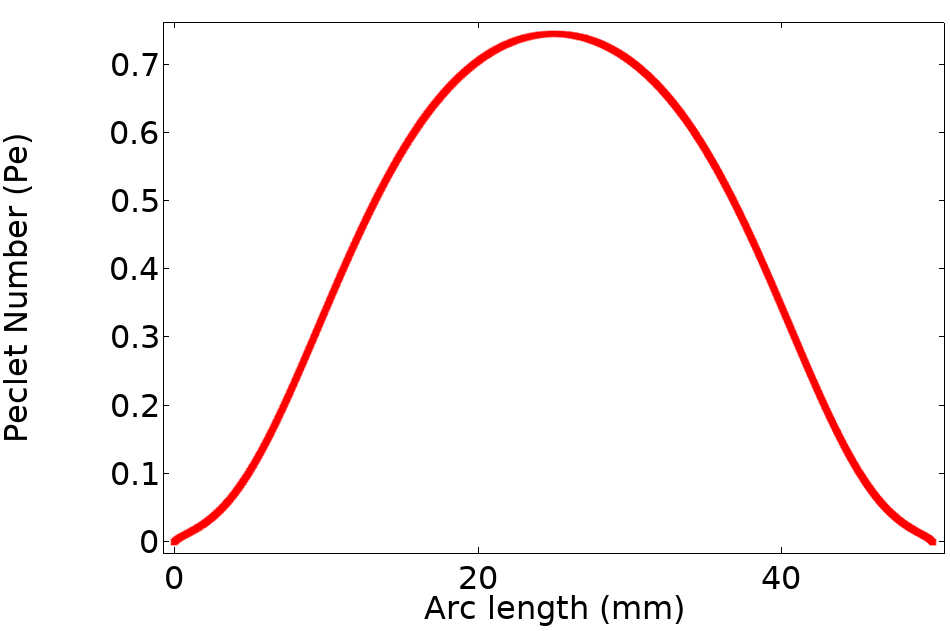

### Shear_profile.png

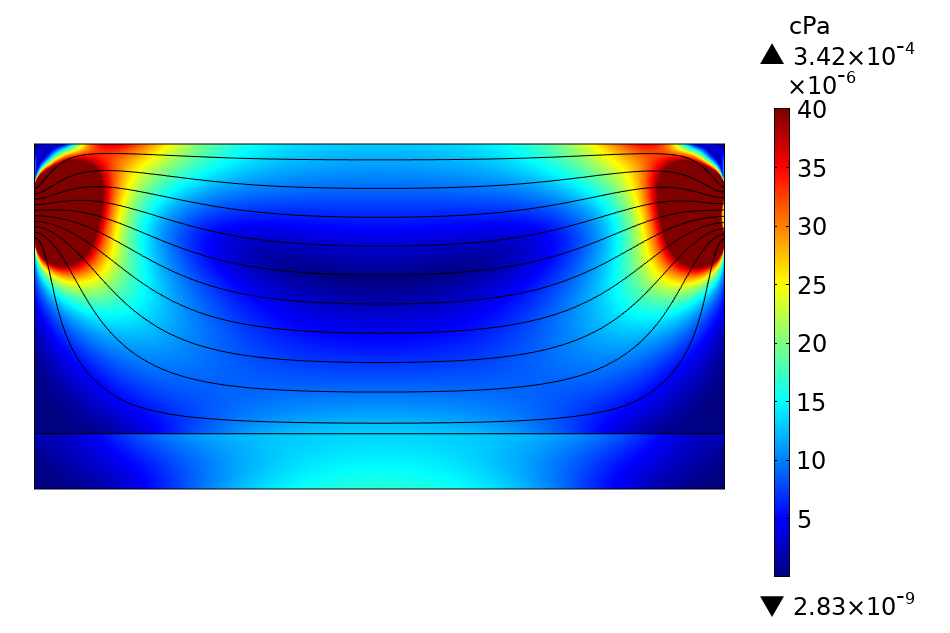

### Shear_profile_scale.png

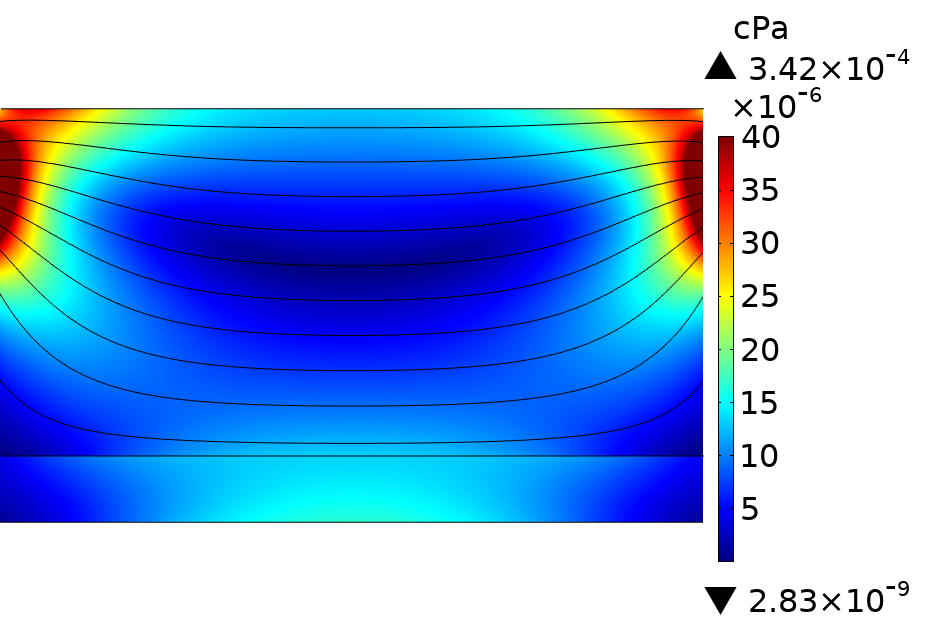

### Velocity_profile.png

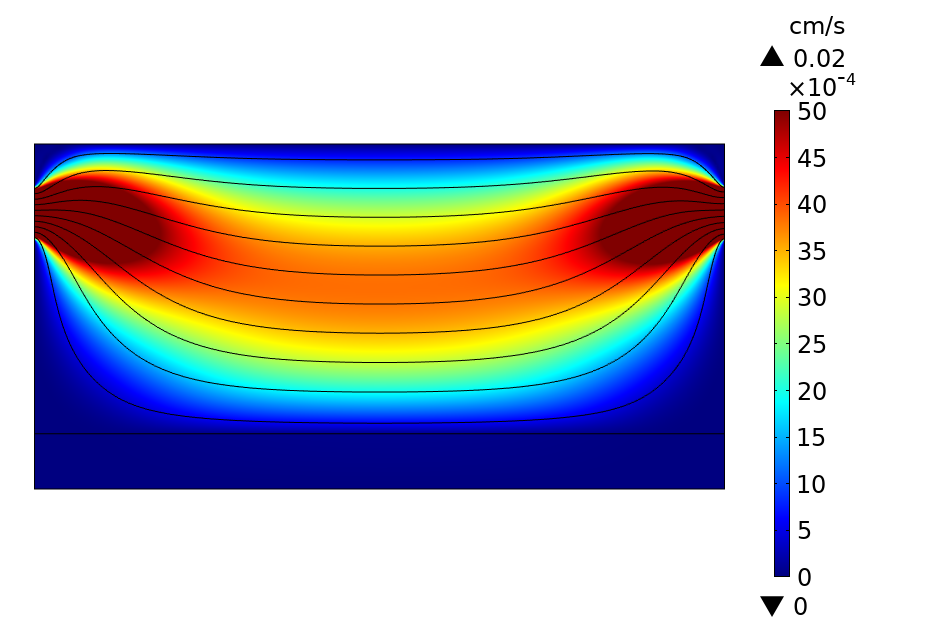

### Velocity_profile_scale.png

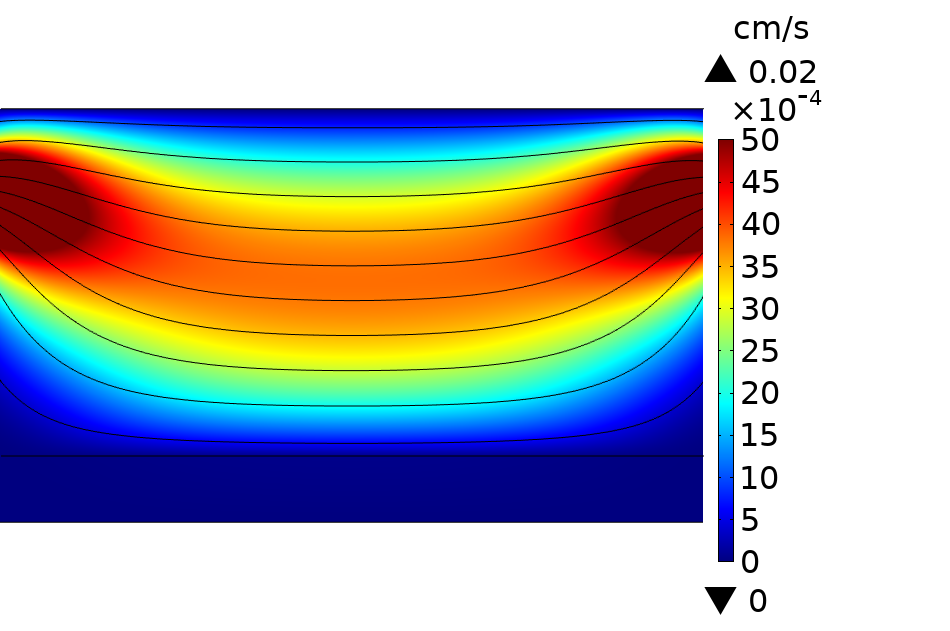
